# Salinity exposure in early-life drives genomic adaptation to climate change in Antarctic toothfish (*Dissostichus mawsoni*)

**DOI:** 10.64898/2026.01.18.700237

**Authors:** Jilda Alicia Caccavo, Enrique Celemín, Pierre de Villemereuil, Marion Gehlen

**Affiliations:** Laboratoire des Sciences du Climat et de l’Environnement, LSCE/IPSL, CEA-CNRS-UVSQ, Université Paris-Saclay, Gif-sur-Yvette, France; Laboratoire d’Océanographie et du Climat Expérimentations et Approches Numériques, LOCEAN/IPSL, UPMC-CNRS-IRD-MNHN, Sorbonne Université, Paris, France; Institute of Biochemistry and Biology, Evolutionary Biology & Systematic Zoology, University of Potsdam, Potsdam, Germany; Institut de Systématique, Évolution, Biodiversité (ISYEB), École Pratique des Hautes Études PSL, MNHN, CNRS, SU, UA, Paris, France; Institut Universitaire de France (IUF); Department of Integrative Biology, University of South Florida, Tampa, FL, United States

**Keywords:** climate genomics, Antarctic toothfish, genotype–environment association, early-life history, fisheries

## Abstract

Antarctic marine ecosystems are undergoing rapid physical change, yet the capacity of long-lived polar fishes to adapt genomically remains poorly understood. Antarctic toothfish (*Dissostichus mawsoni*) are an exploited top fish predator whose early life stages develop beneath the sea-ice edge, where salinity, temperature and circulation are being reshaped by climate change. Using whole-genome resequencing data, we investigated how environmental exposure, particularly during early life, structures patterns of local adaptation in *D. mawsoni*. We analysed 2.4 million unlinked SNPs from 24 adults sampled across the circumpolar distribution of *D mawsoni*, and compared variability in putatively-adaptive loci with variation in environmental parameters from the ORAS5 global ocean reanalysis, including salinity, temperature, mixed-layer depth, sea-ice concentration and thickness, and surface currents. To capture uncertainty in ontogenetic exposure, we constructed three environmental scenarios differing in their spatial and temporal representation of conditions: Point of Capture–Time of Capture (POC-C), Spawning Ground–Time of Birth (SG-B), and Point of Capture–Time of Birth (POC-B). Genotype–environment association (GEA) was performed using redundancy analysis conditioned on fishing pressure and latent-factor mixed models, with high-confidence GEA loci being defined as SNPs jointly detected by both approaches and robust to random-predictor tests. The scenario that best explained genomic variation was SG-B, in which environmental variables were averaged over hypothesised spawning grounds during the egg incubation period of *D. mawsoni* (August - October). Within this scenario, the significant environmental axis, dominated by salinity, was strongly associated with 854 high-confidence GEA loci. Functional enrichment revealed over-representation of gene ontology terms linked to monoatomic ion transport, ion channel complexes and calcium signalling, consistent with salinity-driven selection on osmoregulatory pathways during early development. Our results provide genomic evidence that early-life salinity exposure is a key driver of local adaptation in *D. mawsoni*, underscore the importance of correctly representing life-stage-specific environments in climate genomics, and highlight a concrete pathway by which climate-induced freshening and sea-ice change may alter recruitment and stock resilience.

## 1 Introduction

Despite its status as one of the most pristine environments on the planet, the subzero waters surrounding Antarctica are under threat from compounding anthropogenic influences, including fishing, pollution, tourism, and most critically, climate change (Abram et al., 2025). Heterogenous patterns of climate change impacts across the Southern Ocean, such as on ocean warming, circulation, stratification, acidification, and sea ice loss, can result in divergent repercussions on circumpolarly-distributed species, with extirpation contrasting with population expansions across different regions (Chown et al., 2022; Constable et al., 2023). The rapid progression of climate change impacts, such as recent all-time lows in sea ice extent in the Southern Ocean (Parkinson and DiGirolamo, 2021; Turner et al., 2022), may prevent species from mounting effective responses to environmental change, leading to an accelerated rate of decline and potential extinction (Sage, 2020). This is particularly true for polar species, which are uniquely adapted to the extreme and relatively stable conditions of polar regions (Convey and Peck, 2019), and for whom there exists limited capacity to colonize new habitats (Peck, 2018). Species’ susceptibility to climate change is modulated by their ability to shift their distributions to more suitable habitats, acclimate to new conditions through phenotypic plasticity, or adapt through genomic change (Bernatchez et al., 2024). Analysis of local adaptation in particular can not only reveal to what extent genomic change is contributing to species’ climate change responses, but can also be an indicator of which populations are most at risk across a species’ distribution (Razgour et al., 2019; Barratt et al., 2024). In many cases, adaptive evolution often involves shifts in allele frequencies at many loci rather than the fixation of single beneficial alleles (Bernatchez, 2016). The resulting genomic signatures of local adaptation, i.e., spatial differences at putatively adaptive loci, can be used to identify fish stocks that may be particularly vulnerable to environmental change and thus require a more precautionary management regime (Valenzuela-Quiñonez, 2016; Bernatchez et al., 2017). Species under pressure from fisheries exploitation are particularly vulnerable to climate change impacts (Duncan et al., 2019), with higher risks of isolation, inbreeding, and extirpation (O’Leary et al., 2013; Sadler et al., 2023). Exploitation itself can spur fisheries-induced evolution, resulting in reduced life histories, accelerated maturation, and diminished growth (Heino et al., 2015), making it critical for investigations of local adaptation in exploited species to consider both the influence of climate and fisheries-induced evolution (Waples and Audzijonyte, 2016).

Among exploited species in the Southern Ocean, Antarctic toothfish (*Dissostichus mawsoni*) contribute to the most lucrative fishery, worth over 200 million USD annually (Brooks, 2013; Grilly et al., 2015; CCAMLR Secretariat, 2018, 2019, 2022). *D. mawsoni* are a long-lived, slow-developing top fish predator, reaching up to 2 m in length over a life history during which it takes 12–15 years to achieve sexual maturity, and which can span nearly 30 years (Hanchet et al., 2015; Caccavo et al., 2021). Their life cycle is complex (for review, see Hanchet et al. (2015); La Mesa et al. (2019); Parker et al. (2021); Behrens et al. (2021); Di Blasi et al. (2024)), resulting in Southern Ocean distributions that range from nearshore continental shelf areas to deeper slope and offshore sea mounts. Two aspects of *D. mawsoni* life history render them particularly vulnerable to climate change impacts: 1) early-life association with sea ice (Parker et al., 2021); and 2) dependence on current systems (Ashford et al., 2017). Their sedentary life strategy, common across Antarctic fish species (Zimmermann and Hubold, 1998), necessitates a reliance on ocean circulation to complete their life cycle and exchange individuals between regional populations (Caccavo et al., 2021). Eggs hatch underneath sea ice near its offshore edge during the austral spring, where the unique surface environment provides both protection as well as access to abundant phytoplankton food sources (Parker et al., 2021). Models project that climate change impacts will intensify in the Southern Ocean over the course of the 21*^st^* century (Masson-Delmotte et al., 2021), with substantial changes in sea ice extent that are consistent with the record sea ice lows recorded in recent years (Parkinson and DiGirolamo, 2021; Turner et al., 2022). Furthermore, a changing climate drives shifts in circulation patterns, altering the position and intensity of currents, with grave consequences on fish life history connectivity (Petitgas et al., 2013; Wilson et al., 2016; Young et al., 2018; IPCC, 2019; Dell’Apa et al., 2023). Such conditions foster selection for individuals able to resist changing conditions in early life, giving rise to the hypothesis that climate-driven selection is acting on early life stages of *D. mawsoni*. Indeed, early life stages often serve as bottlenecks for selection due to their heightened vulnerability to environmental variability (Pörtner and Farrell, 2008; Alix et al., 2020; Chen and Liu, 2022). Consequently, early life stages (i.e., eggs and larvae) of *D. mawsoni* are the most at-risk from climate change and the most likely to exhibit adaptation to environmental shifts, in part because of their restricted ability to shift their distribution in response to environmental change (Llopiz et al., 2014).

Population genetic structure in *D. mawsoni* is characterized by high levels of gene flow throughout the Southern Ocean, leading to a general state of panmixia across regional populations (Mugue et al., 2014; Ceballos et al., 2021; Choi et al., 2021; Maschette et al., 2023; Caccavo et al., 2025). However, we recently showed that this pattern of gene flow is susceptible to interruption over time, with drops in connectivity between cohorts ascribed to variable recruitment (i.e., the proportion of fish reproducing in a given year), which may ultimately be driven by fisheries pressure and climate change (Caccavo et al., 2025). Indeed, a growing body of research demonstrates that even in marine environments, where reduced geographic barriers to gene flow often support panmictic populations, adaptive divergence can arise (Gleason and Burton, 2016; Attard et al., 2018; Celemín et al., 2023; Klein et al., 2024). In our recent analysis of population dynamics in *D. mawsoni* based on whole-genome resequencing (WGR) data, we discovered over 100 thousand single-nucleotide polymorphism (SNP) outliers based on population structure (Luu et al., 2017), which represent potential targets of selection acting on the toothfish genome (Caccavo et al., 2025). Thus, this dataset from Caccavo et al. (2025), with individuals from across the circumpolar distributions of *D. mawsoni*, including all fisheries management areas, permits us to investigate the extent to which climate change may be driving local adaptation in toothfish.

In this study, we employed a genotype-environment association (GEA) analysis based on multiple approaches to increase confidence in the putatively adaptive loci identified (Lotterhos, 2023): 1) a multivariate approach through redundancy (RDA) analysis (Capblancq et al., 2021); 2) a univariate approach, through latent-factor mixed modelling (LFMM) (Frichot et al., 2015); and 3) removal of false positives (Salloum et al., 2022). In our analysis, we carefully consider how we represent environmental exposure in *D. mawsoni*, given the key role that we hypothesize early life exposure to play in driving local adaptation Our primary goal was to determine which environmental variables best explain genomic variation in *D. mawsoni*, and to refine the set of candidate SNPs associated with local adaptation.

## 2 Materials and Methods

### 2.1 Genomic data

The genomic data used in this study derive from a circumpolar set of 24 samples of *D. mawsoni* (Figure 1, Table S1). A detailed description of the preparation of single-nucleotide polymorphism (SNP) libraries can be found in Caccavo et al. (2024), including DNA extraction from *D. mawsoni* tissue and otoliths, preparation of whole-genome libraries, sequencing, bioinformatics processing, and SNP calling. These same genomic data were used in Caccavo et al. (2025) to confirm panmixia across the circumpolar distribution of *D. mawsoni*, while highlighting reductions in gene flow linked to fisheries pressure and cohort. Indeed, the relatively low levels of structure identified across *D. mawsoni* populations reduce the risk of identifying false positives in genome scans for local adaptation (Salloum et al., 2022). In short, we used *mpileup* in BCFTOOLS v1.17 (Danecek et al., 2021) to call genotypes from fully processed reads downsampled to 5× coverage (for details, see Caccavo et al. (2024) and Caccavo et al. (2025)). SNPs were filtered based on minimum base pair quality (`QUAL < 20`), mapping quality (`MQ < 30`), minor allele frequency (`MAF < 0.05`), allowing only for SNPs present in at least 75% of individuals (`AN/2 < 0.75n`), removing non-biallelic SNPs (`--max-alleles 2`), and excluding indels (`--exclude types indels`). To identify and filter linked SNPs, we used PLINK2 `--indep-pairwise`, applying a 20 kb cutoff and minimum weight setting of 0.5, resulting in 2.8 million filtered SNPs, of which 2.4 million were unlinked. This set of 2.4 million unlinked SNPs was used in the genotype-environment association (GEA) analysis.

**Figure 1.**
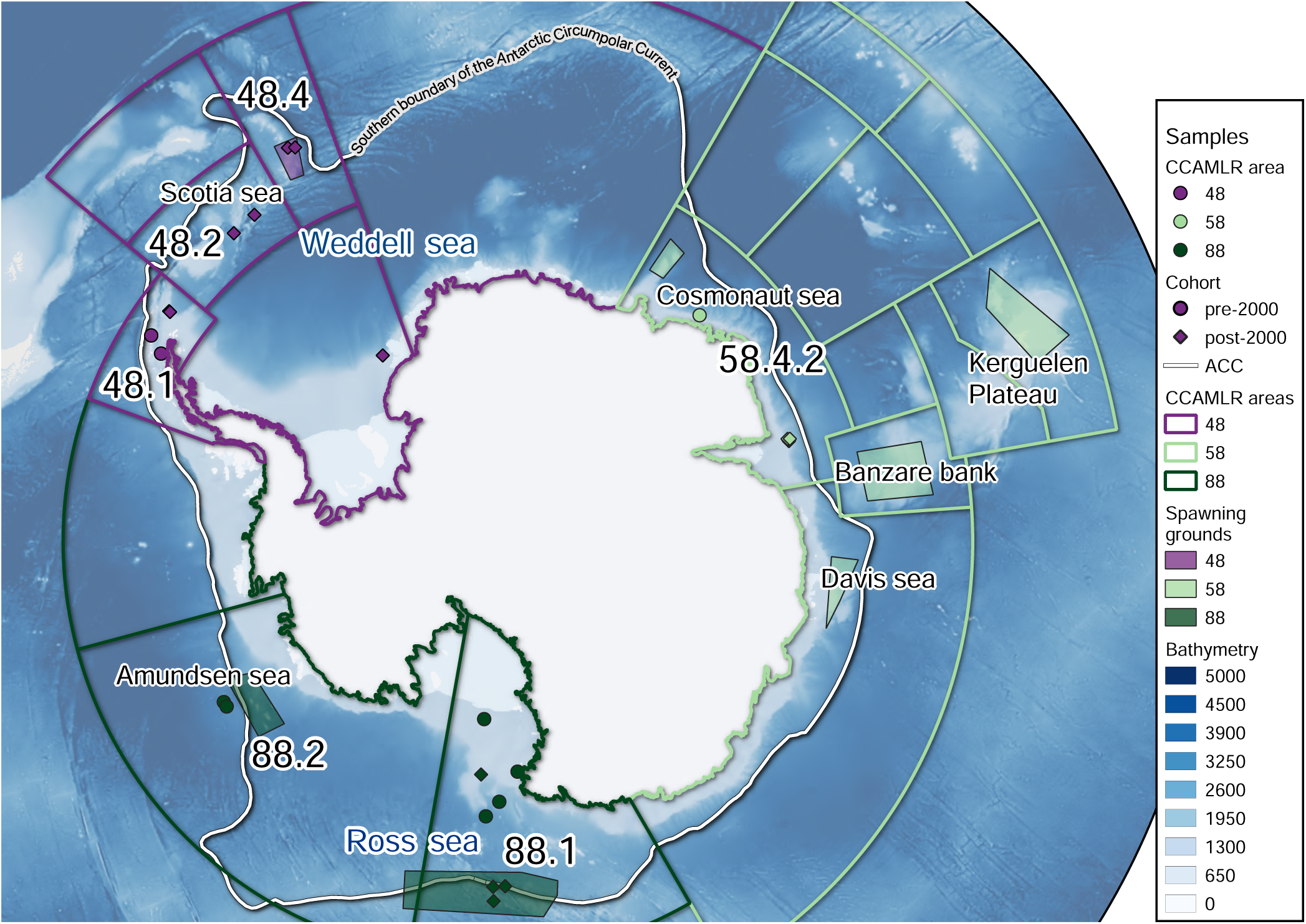
Map showing the provenance of *D. mawsoni* samples, including spawning grounds based on Dunn et al. (2012).

### 2.2 Environmental representation and data

#### 2.2.1 Environmental scenarios

We were obliged to carefully consider how we represent environmental exposure in *D. mawsoni*, given the key role that we hypothesize early life exposure to play in driving local adaptation. Indeed, we can consider the role of environment in driving selection pressure across three different scenarios which vary across uncertainty (i.e., the likelihood that the values of environmental variables used are representative of the environmental exposure that the fish actually experienced), grounding in life history, time, and space (Table 1). The first scenario, which we refer to as the Point of Capture–Time of Capture (POC-C) scenario, posits that the location where fish were captured represents the area where environmental exposure exercised the greatest impact on selection pressure, at the time when the fish were captured. This first scenario, which we can consider as the null hypothesis, has the lowest level of uncertainty, but also the lowest grounding in our understanding of *D. mawsoni* life history.

**Table 1.** Environmental exposure scenarios Point of Capture–Time of Capture (POC-C), Spawning Ground–Time of Birth (SG-B), Point of Capture–Time of Birth (POC-B), and the extent to which they fulfil criteria related to uncertainty (i.e., the likelihood that the values of environmental variables used are representative of the environmental exposure that the fish actually experienced), and grounding in life history, as well as in our understanding of the timing of when climate change impacts are likely to exert the greatest impact on selection (i.e., during early life history).

|  | POC-C | SG-B | POC-B |
| --- | --- | --- | --- |
| Uncertainty | Low | High | Low |
| Life history | Low | High | Low |
| Timing (early) | Low | High | High |

The second scenario, or alternative hypothesis, has the highest level of uncertainty, but the greatest grounding in *D. mawsoni* life history. In this second scenario, referred to as Spawning Ground–Time of Birth (SG-B), we propose that the environmental conditions at the location where fish were born (the hypothesized spawning ground), at the estimated time of birth, were the greatest drivers of selection pressure. While the SG-B scenario is grounded in our understanding of the critical role of early life history in driving climate change adaptation in fish (Chen and Liu, 2022), there are two uncertainties associated with environmental representation in this dataset. First, the only confirmed spawning ground for *D. mawsoni* is above offshore sea mounts in the Ross Sea (Parker et al., 2019, 2021); the rest are hypothesized based on similar conditions found in parts of other circumpolar habitats of *D. mawsoni*, as well as distributions of gravid fish in these areas (Dunn et al., 2012). Second, no age data were available for the *D. mawsoni* samples used in this study. This means that we had to rely on total length data to estimate birth years based on age-length relationships derived from *D. mawsoni* von Bertanlaffy growth curves (Horn, 2002; Brooks et al., 2011). The issue with this approach is that after fish reach sexual maturity at ∼15 years of age, the von Bertanlaffy curves begin to flatten out, making it difficult to estimate whether a fish of a given length is e.g., 15, 20, or 25 years old. Luckily for this study, we had fish caught during two separate catch periods: the early 2000s, and the 2010s. This allowed us to clearly distinguish between early life exposure of fish caught in the early 2000s, which we estimated to be in the mid-1980s, and fish caught in the 2010s, which we estimated to be at various points during the 2000s.

Finally, the third scenario represents a middle ground between the POC-C and the SG-B scenarios. This third scenario, Point of Capture–Time of Birth (POC-B), retains the hypothesis from SG-B that environmental exposure during early life has the greatest impact on selection pressure, but reduces uncertainty by deriving environmental exposure data from the location where fish were captured, rather than hypothesized spawning ground locations. However, there is little evidence that fish remain in the same location throughout their life history, thus reducing the extent to which this POC-B scenario is grounded in our understanding of *D. mawsoni* life history. Indeed, while tagging studies reveal that *D. mawsoni* largely engage in a sedentary life strategy, there is nonetheless evidence that a small proportion engage in long-distance migrations (Grilly et al., 2022). Furthermore, evidence from age class distributions (Hanchet et al., 2008), spawning grounds (Parker et al., 2019, 2021), and biophysical interactions (Ashford et al., 2012) support a more complex life history hypothesis for *D. mawsoni* (Hanchet et al., 2015). This hypothesis posits that fish largely do not remain in the same location throughout their life, but rather, engage in life history migrations driven by gyre circulation (Ashford et al., 2017). After spawning takes place on offshore sea mounts (Hanchet et al., 2008), eggs and larvae develop near the surface under the sea ice edge (Parker et al., 2021). As early life stages develop, they are carried with gyre circulation towards the continental shelf (Behrens et al., 2021), where early juveniles settle in benthopelagic habitats, gaining condition on nutrient rich prey (Hanchet et al., 2008; La Mesa et al., 2019). Maturing fish then continue with gyre circulation away from the continental shelf and back towards offshore sea mounts to spawn (Parker et al., 2019; Di Blasi et al., 2024), hence closing the recruitment loop.

#### 2.2.2 Environmental data

Using the capture year and total length of samples, we derived 4 estimated birth years (1985, 2000, 2005, and 2010) across two cohorts (fish born before 2000 and fish born after 2000). Across all scenarios, we aimed to represent five environmental parameters that are both critical to *D. mawsoni* life history, and indicative of climate change impacts in the Southern Ocean: 1) sea ice; 2) primary productivity; 3) salinity; 4) temperature; and 5) current strength. To represent these five parameters, we used 8 environmental variables: 1) sea ice thickness (m); 2) sea ice concentration; 3) mixed-layer depth at 0.01 density (m); 4) mixed-layer depth at 0.03 density (m); 5) salinity (PSU); 6) sea surface temperature (°C); 7) meridional current velocity (m/s); and 8) zonal current velocity (m/s) (Table S2). Because *D. mawsoni* spawn between August and October, we extracted monthly averages for variables during August, September and October across the four estimated years. For the POC-C dataset, we also used monthly averages for variables from August through October to be consistent with the Time of Birth (SG-B and POC-B) scenarios, but instead derived these data from capture years. Environmental exposure data in the SG-B scenario was based on the average value for each variable over the area of the hypothesized spawning ground assigned to each individual fish. Fish were assigned spawning grounds based on their capture location and the location of hypothesized spawning grounds as defined in Dunn et al. (2012) (Table S3). We could not rely on observational data to extract these variables due to its spatial patchiness in the Southern Ocean, particularly for the pre-2000 cohort in the Time of Birth scenarios. For this we reason, we used the ORAS5 global ocean reanalysis that combines model data and observations to create a spatially and temporally complete dataset of monthly averages from 1958 to present at a resolution of 0.25° x 0.25° (equivalent to approximately 10 km in the Southern Ocean) (CMEMS, 2021). It is possible to observe clear spatial variation in all eight environmental variables across all four birth years (Figure S1 through Figure S8), which emphasizes the importance of considering environmental exposure at the appropriate time during which fish were in their early life stages. Indeed, environmental parameters vary considerably across scenarios themselves (Figure 2), with SG-B typically displaying greater variability due to the larger area occupied by the ensemble of confirmed and proposed *D. mawsoni* spawning grounds (Figure 1, Table S3).

**Figure 2.**
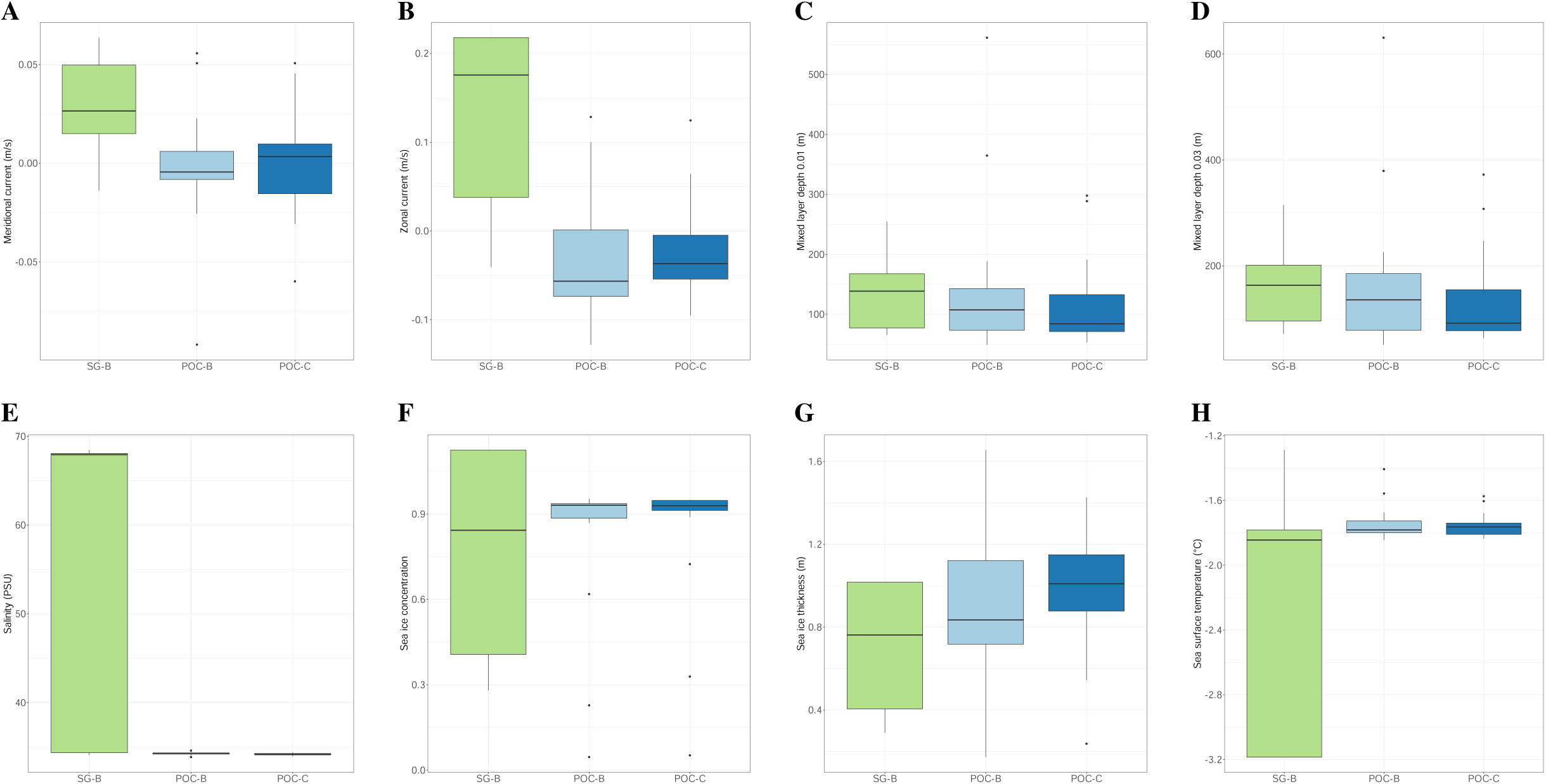
Environmental variability among locations represented in scenarios Point of Capture–Time of Capture (POC-C), Spawning Ground–Time of Birth (SG-B), and Point of Capture–Time of Birth (POC-B), across (A) meridional current velocity (m/s), (B) zonal current velocity (m/s), (C) mixed layer depth density 0.01 (m), (D) mixed layer depth density 0.03 (m), (E) salinity (PSU), (F) sea surface temperature (°C), (G) sea ice concentration, and (H) sea ice thickness (m).

Predictably, strong correlations existed between many environmental variables across all three scenarios (Figure S9). When running a principal component analysis (PCA) among variables in each scenario, we discovered that the first three principal components (PCs) explained most of the variance among environmental variables across all three scenarios (SG-B: 98.2%, POC-B: 80.1%, POC-C: 89.0%) (Figure S10). Across scenarios, salinity, sea ice, and current were the environmental parameters most responsible for driving the variation explained by the first three PCs, with the PCs within the POC-B and POC-C scenarios also being driven by primary productivity (Figure S11). Because GEA requires that collinearity between environmental variables be constrained (i.e., *r ≤ |*0.7|) (Frichot et al., 2015; Capblancq et al., 2021), we opted to use the PC scores for the analysis, derived from the first three PCs of the PCA run in each scenario, rather than use the highly correlated raw environmental data (Table S4).

### 2.3 Fishing data

For exploited species, fishing pressure itself may induce genomic adaptation, resulting in changes in growth rates, age-at-maturity, and maximum size(Heino et al., 2015). For this reason, we included fishing data in our analysis, provided by the Commission for the Conservation of Antarctic Marine Living Resources (CCAMLR) through a formal C2 data request, including the following information for each haul set at the locations and time periods requested: 1) hook count; 2) fish greenweight; 3) individual fish count; 4) catch per unit effort (CPUE). For the estimated birth years within the SG-B and POC-B scenarios (1985, 2000, 2005, and 2010), no fishing was recorded in 1985, and within management area 48, no fishing took place during any birth year. While collinearity was expectedly high between fishing data types (Figure S12), we ultimately chose to include only individual fish count in the GEA, as this was the data type most likely to represent the genomic impact of fishing pressure based on the removal of individuals from populations due to fishing (Table S4).

### 2.4 Genotype-environment association

To perform the GEA analysis, we employed a multivariate (redundancy analysis, RDA), and a univariate (latent factor mixed modelling, LFMM) approach. This two-pronged approach improves the robustness of association detection by optimizing the trade-off between false negatives and false positives, thereby reducing the likelihood of missing relevant SNPs and emphasizing loci consistently identified across methods (Lotterhos, 2023). To further enhance our confidence in the detected loci, we also performed a false-positive analysis using both RDA and LFMM to associate our genomic data with randomly generated environmental data (Salloum et al., 2022). Any SNPs identified that met association criteria in either method were removed from the final SNP set.

#### 2.4.1 Redundancy analysis

Unlinked SNP genotypes stored in *vcf* format were first converted to *raw* format using PLINK2 for analyses in R using the package *vegan* (Oksanen et al., 2019). We then read in the *raw* file containing the 2.4 million unlinked SNPs from the 24-individual circumpolar *D. mawsoni* dataset using `read.PLINK`. After checking for and imputing missing SNPs, we performed a forward selection using the function `forward.sel` from the R package *adespatial* (Dray et al., 2025), adjusting resultant *p*-values based on Holm correction for multiple tests. RDA models were only developed for scenarios (POC-C, SG-B, or POC-B) for which significant predictor variables were identified in the forward selection analysis. Because no significant genetic structure was observed previously in the dataset (Caccavo et al., 2025), we did not include spatial variables as a condition in the RDA. Instead, we used the individual counts from the fishing data as a conditioning variable to distinguish between adaptation linked to environmental parameters and adaptation associated with fisheries-induced evolution (Heino et al., 2015). After confirming that there was no collinearity between the three PCs derived from the raw environmental data (*r ≥ |*0.7|), we ran a constrained RDA (pRDA) with the PC scores from the first three PCs as the predictor variables and the individual counts from fishing data as the conditioning variable using the function `rda` (Capblancq et al., 2021). Variance inflation factor (VIF) values were checked to assure that no component of the pRDA had a VIF > 3. We checked the significance of the predictor variables by running an analysis of variance (ANOVA) with 1000 permutations using the function `anova.cca`. We used a cut-off of three standard deviations (*p* = 0.0027) to identify putatively adaptive SNPs associated with significant predictors based on SNP loading scores from the first two RDA axes (Forester et al., 2018).

#### 2.4.2 Latent factor mixed modelling

We associated imputed unlinked SNP genotypes to each set of PC scores separately using LFMM as implemented in the R package *lfmm* (Caye et al., 2019). To identify the appropriate number of K latent factors for the LFMM analysis, we used the function `snmf` as implemented in the R package *LEA* (Frichot et al., 2015) for K = 1 - 7, choosing the K value that produced the lowest cross-entropy value when assessing the `snmf` output using the function `cross.entropy`. This led to the selection of K = 1 latent factors for the subsequent LFMM analyses based on having the lowest cross-entropy value. We then ran the LFMM analyses using K = 1 latent factors for each set of PC scores using the function `lfmm_ridge`, deriving *p*-values with the function `lfmm_test`. To account for multiple testing, we applied a false discovery rate of *q* < 0.1 with the function `qvalue` from the R package *qvalue* (Storey et al., 2025), retaining only SNPs that met this criteria.

#### 2.4.3 Selection of high-confidence GEA loci

We re-ran the RDA and LFMM analyses with the same parameters but now using 100 sets of random predictor variables to identify false positive loci that generated bogus associations. Values were randomly generated from a normal distribution using the base R function `rnorm` for *n* = 24 values with a mean of 0 and a standard deviation of 1. Loci that generated significant associations in more than 5% of the random predictor tests (i.e., loci associated with more than five random predictors) were considered false positives and removed from the analysis. Putatively-adaptive SNPs that were significantly associated with predictor variables in both the RDA and LFMM analyses and did not associate with more than five random predictors were considered high-confidence GEA loci.

### 2.5 Gene ontology

We extracted gene ontology (GO) terms from our set of high-confidence GEA loci to perform a functional enrichment analysis based on the annotated genome of the closely related *Dissostichus eleginoides* (Lee et al., 2024). To do this, we first obtained sequences associated with each high-confidence GEA locus by extracting 300-bp regions flanking either side of each locus (i.e., 601-bp sequence per locus) using the tool `getfasta` in BED_TOOLS V_2.31.1 (Quinlan and Hall, 2010). We then used BLASTN to retrieve gene functions from the *D. eleginoides* genome using the 601-bp sequences associated with each high-confidence GEA locus position in the *D. mawsoni* genome. Retrieved sequences were filtered based on sequence length and *E*-value (i.e., the number of expected hits of similar quality that could occur by chance), retaining only sequences within the range of 550 - 650 bp where *E* < 1e*^−^*^50^ and identity >95%, such that no more than one gene per high-confidence GEA locus would be retained. We used MAGMA (de Leeuw et al., 2015) to locate genes within 20 kb upstream or downstream from high-confidence GEA loci. To prepare the input for the GO term analysis in DAVID (Sherman et al., 2022), we used KOBAS (Bu et al., 2021) to retrieve official gene symbols from the zebrafish (*Danrio rerio*) genome annotation based on our retrieved FASTA sequences. We submitted our gene list from KOBAS into DAVID, selecting OFFICAL_GENE_SYMBOL as our identifier and *D. rerio* as our species. Using the Functional Annotation tool, we expanded the GO to reveal terms associated with molecular functions (MF), cellular components (CC), and biological processes (BP). We exported fold enrichment values, and used false discovery rate (FDR) post-hoc tests to derive *p*-values for GO terms; only GO terms with *p* < 0.01 were considered significant.

## 3 Results

### 3.1 Genotype-environment association

Forward selection identified PC1 (largely informed by salinity, see Figure S11a) as significant in the Spawning Ground– Time of Birth (SG-B) scenario with an adjusted *p*_PC1_ = 0.018 (Table 2). No PCs were identified as significant in the forward selection for the Point of Capture–Time of Birth (POC-B) scenario, and while PCs 1 and 3 were initially identified in the forward selection for the Point of Capture–Time of Capture (POC-C) scenario, they fell short of significance after multiple test correction (adjusted *p*_PC1_ = 0.162, adjusted *p*_PC3_ = 0.323) (Table 2). RDA model development and candidate SNP selection were thus only carried out using predictor variables from the SG-B scenario. The RDA model constrained by fishing pressure (pRDA) based on data from the SG-B scenario revealed significant associations between genomic variation and PC1 (*F*_1,19_ = 1.2645, *p*_PC1_ = 0.002) (Table 3). The pRDA biplot reveals the extent to which genomic responses to predictor variables are partitioned among CCAMLR fisheries management areas, with PC1 (salinity) driving the separation between cohorts in area 88, and to a lesser extent between areas 48 and 58 (Figure 3). Among the 2.4 million unlinked SNPs used in the analysis, 17,928 were identified as putatively adaptive associated with PC1 (salinity) based on the SG-B pRDA model, while the univariate latent factor mixed modelling (LFMM) approach revealed 12,562 putatively-adaptive SNPs. False positive analysis identified 0 spurious SNPs using the pRDA model, which could be linked to the lack of genetic structure in the *D. mawsoni* dataset (Salloum et al., 2022; Caccavo et al., 2025). While false positive analysis using the LFMM approach identified 15,190 spurious SNPs, only 4,692 of these were among the 12,562 putatively-adaptive SNPs originally identified in the LFMM analysis. Thus, 7,870 putatively-adaptive SNPS were retained after removal of false positives from the LFMM dataset. Finally, 854 putatively-adaptive SNPs shared between both the pRDA and LFMM datasets were retained as high-confidence GEA loci for the gene ontology (GO) analysis. These results are summarized in Table 4.

**Figure 3.**
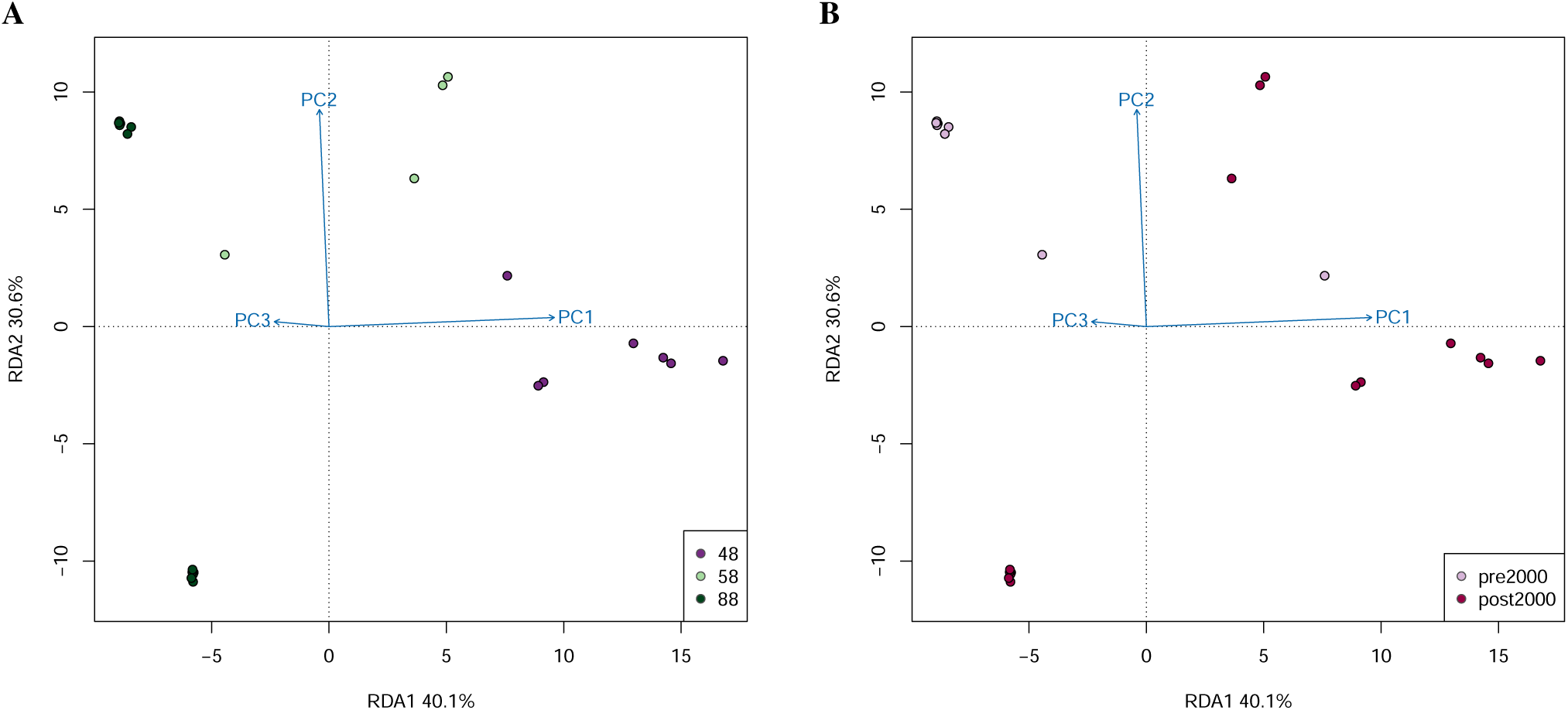
Biplot of the RDA model constrained by fishing pressure (pRDA) from the Spawning Ground–Time of Birth (SG-B) scenario, coloured by (A) CCAMLR fisheries management area (48, 58, and 88), and (B) cohort (pre-2000, post-2000). Predictor variables derived from principal components (PCs) 1–3 represented by blue arrows, with PC1 largely driven by salinity, PC2 by sea ice, and PC3 by current (Figure S11a). Note that only PC1 was identified as significant.

**Table 2.** Forward selection analysis based on the predictor variables PC scores 1–3 in each scenario: Point of Capture–Time of Capture (POC-C), Spawning Ground–Time of Birth (SG-B), and Point of Capture–Time of Birth (POC-B). Only significant results are shown, noting that the PCs initially identified as significant in the POC-C scenario were ultimately not significant after correction for multiple tests (adjusted *p*-value).

| Scenario | Predictor variable | R2 | R2 Cumulative | AdjR2Cum | F | p-value | Adjusted p-value |
| --- | --- | --- | --- | --- | --- | --- | --- |
| SG-B | PC1 | 0.05148626 | 0.05148626 | 0.00837200 | 1.19418200 | 0.001 | 0.018 |
| POC-C | PC1 | 0.04856934 | 0.04856934 | 0.00532249 | 1.12307200 | 0.009 | 0.162 |
| POC-C | PC3 | 0.04824548 | 0.09681481 | 0.01079718 | 1.12175800 | 0.019 | 0.323 |

**Table 3.** F- and *p*-values from the analysis of variance (ANOVA) assessing the significance of predictor variables (principal component (PC) scores 1–3) in the pRDA model from the scenario Spawning Ground–Time of Birth (SG-B). The percent variance explained by each predictor variable in the pRDA model is also shown.

| Predictor variable | F-value | p-value | % Variance explained |
| --- | --- | --- | --- |
| PC1 | 1.2645 | 0.002 | 40.1 |
| PC2 | 0.9788 | 0.596 | 30.6 |
| PC3 | 0.9528 | 0.898 | 29.2 |

**Table 4.**
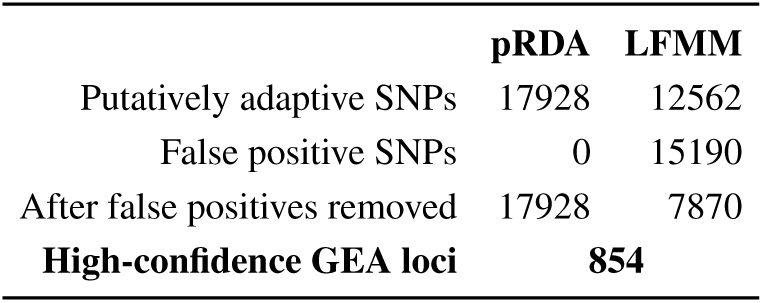
High-confidence GEA loci derived from putatively adaptive SNPs identified in both the multivariate RDA model constrained by fishing pressure (pRDA) and the latent factor mixed modelling (LFMM) univariate approach after removal of false positives.

### 3.2 Gene ontology

Through functional enrichment analyses we found 23 GO terms across molecular function, cellular component, and biological process categories that were significantly (*p* < 0.01) overrepresented among high-confidence GEA loci with respect to the whole genome, many of which play a role in osmoregulation (Figure 4, Table S5). With respect to the molecular function category (*n* = 9), we identified GO terms related to calcium (11 genes), voltage-gated calcium (8 genes), and monoatomic ion (21 genes) channel activity, as well as calcium ion binding (57 genes) (Table S6). Among cellular component GO terms (*n* = 5), we identified 16 genes related to the monoatomic ion channel complex, which are essential for electrical signalling, osmotic balance, and many forms of cell communication (Hoffmann et al., 2009; Eugenin et al., 2022). Finally in the biological process category (*n* = 9), we observe once again an over-representation of ion transport-related genes: calcium ion transmembrane (12 genes), monoatomic ion (42 genes), calcium ion (12 genes), and monoatomic ion transmembrane (30 genes) transport.

**Figure 4.**
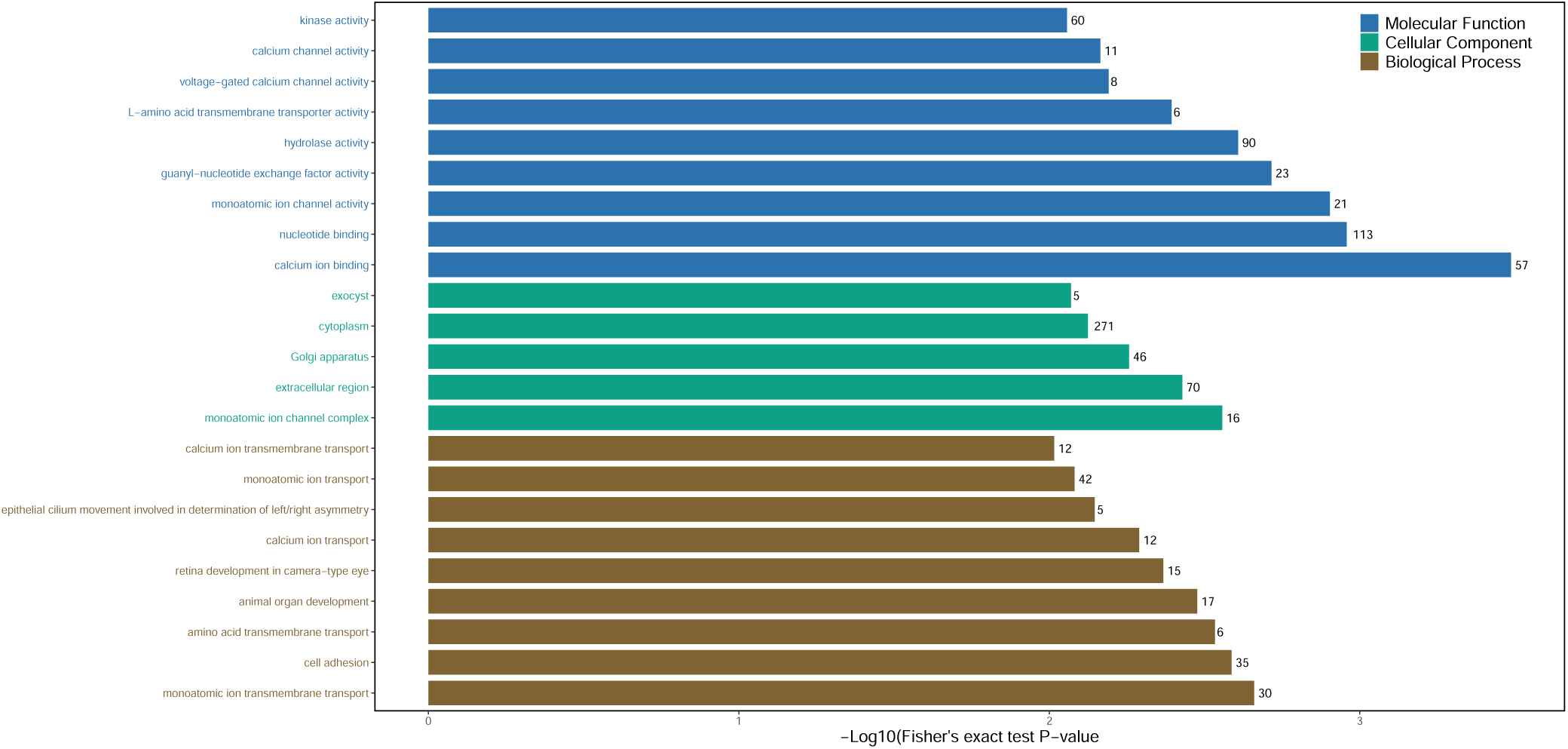
Gene ontology (GO) terms significantly (*p* > 0.01) overrepresented with respect to the whole genome, derived from high-confidence GEA loci associated to predictor variable PC1 (largely driven by salinity, Figure S11a) from the Spawning Ground–Time of Birth (SG-B) scenario. GO terms are coloured by category: molecular function (blue), cellular component (green), or biological process (brown). Y-axis labels refer to the GO terms listed in Table S5. Bar lengths correspond to the log transformed *p*-value of each GO term. At the end of each bar, the number of genes identified within the GO term is indicated (Table S6).

## 4 Discussion

### 4.1 The impact of environmental exposure scenarios in climate genomics

Among the three environmental scenarios that we investigated, only one, the Spawning Ground–Time of Birth (SG-B) scenario, revealed significant associations with the principal component (PC) scores derived from our set of eight environmental variables, in particular the PC most informed by salinity (Figure S11a). This scenario was most well-grounded in our understanding of *D. mawsoni* life history, though it is limited by uncertainties in the exact locations and years during which the fish analysed in our study were experiencing their most early life stages, i.e., egg incubation.

There is growing recognition that genotype-environment association (GEA) results are highly sensitive to how the environment is represented, i.e., which datasets and variables are used, their spatial and temporal resolution, and the life stage or exposure window they are intended to capture (Hoban et al., 2016; Flanagan et al., 2018; Dauphin et al., 2023). Recent empirical work shows that changing the spatial resolution or type of environmental proxies can substantially alter which loci are identified as associated with environment (Barratt et al., 2024; Guillaume et al., 2024), underscoring the need to align environmental predictors with the ecology and ontogeny of the focal species.

Our results lead us to hypothesize that future analyses based on datasets of fish of known age, paired with a greater understanding of circumpolar *D. mawsoni* spawning grounds, would yield even larger effect sizes, with a potentially greater number of PCs identified as significant (i.e., PCs 2 and 3, associated with sea ice and current, respectively). Lacking age data, a dataset based on immature fish (i.e., mid- and late-stage juveniles 10–15 years of age) would be associated with reduced uncertainty surrounding the estimation of birth years, given the more linear manner with which *D. mawsoni* grow prior to reaching sexual maturity (Horn, 2002; Brooks et al., 2011). However, juveniles are rarely caught by the fishing industry (Hanchet et al., 2008), and thus would require procurement via scientific surveys, which ultimately produce far fewer samples, and typically have a regional focus, making circumpolar comparisons across heterogeneous climate change regimes difficult. While it remains a perennial necessity to compromise between the best available samples and the best possible study design, such constraints and aspirations are critical to consider when devising future climate genomics studies of long-lived marine species like *D. mawsoni*.

### 4.2 The role of salinity in driving genomic change

Sea ice formation and melt can strongly influence salinity, due to the processes by which salt brine is expelled from sea water upon freezing, and by which meltwater can freshen the surface layer (Willis et al., 2023). A clear question that arises from our analysis is why, if sea ice is the driver of salinity changes, did we not observe a direct association between genomic adaptation and sea ice? Indeed, the two environmental variables that we used to describe the parameter sea ice, sea ice thickness and sea ice concentration, were both partitioned within PC2 (Figure S11a), which was not at all found to be associated with variation in our putatively adaptive loci (Table S6). However, if we consider the role of salinity in the development of fish eggs generally, as well as *D. mawsoni* eggs specifically, a more comprehensible explanation for our results arises. Indeed, while plankton buoyancy is typically determined by both temperature and salinity, buoyancy in fish eggs in particular is governed principally by salinity (Sundby and Kristiansen, 2015). Sea ice formation increases the salinity and thus the density of underlying shelf waters through brine rejection, enhancing vertical mixing and overturning circulation (i.e., reducing stratification) as dense, high-salinity water descends to the ocean floor (Meredith et al., 2017). In regions where winter sea ice formation is an important source of dense water, a reduction in sea-ice extent or duration is expected to freshen the surface layer, reducing water density and increasing stratification (Komatsu et al., 2025). A key aspect of *D. mawsoni* egg development is that they float from where they are spawned near the seafloor above offshore sea mounts, to the surface below the sea ice edge (Parker et al., 2021). This requires that the buoyancy of *D. mawsoni* eggs be below the density of the surrounding sea water, in order to float to the surface. Loss of sea ice would freshen the surrounding waters, reducing salinity, which would in turn decrease surface water density and increase stratification, ultimately rendering it more difficult for *D. mawsoni* eggs to float to the surface, where the under ice environment ultimately provides protection and critical food sources for developing larvae (Parker et al., 2021). Beyond impacts on buoyancy, salinity can also influence the development and survival of fish eggs, impacting egg hydration and perivitelline fluid composition, ultimately leading to reduced hatching and higher embryo mortality (Bœuf and Payan, 2001). While few studies exist investigating the impact of changing salinity on Antarctic fish (Mintenbeck et al., 2012), a handful of physiology studies on coastal, benthic species in the Southern Ocean have revealed a tendency towards worse outcomes for fish exposed both to reduced salinity and increasing temperatures (Rodrigues et al., 2013; Navarro et al., 2019; Martínez et al., 2021; Vargas-Chacoff et al., 2021).

Until 2014, in contrast to the Arctic, most regions of the Southern Ocean were experiencing increases in sea ice extents, with the main exceptions being the Amundsen and Bellingshausen Seas, as well as certain parts of the Weddell Sea, where consistent sea ice losses over time were observed (Parkinson, 2019). Recently however, the Southern Ocean has witnessed record-low sea ice extents, driven by strong negative anomalies in several Southern Ocean sectors where *D. mawsoni* occur (e.g., the Ross and Weddell Seas) (Parkinson and DiGirolamo, 2021; Turner et al., 2022). These trends are reflected in the data used for our analysis from the ORAS5 global ocean reanalysis, with pre-2000 cohorts of *D. mawsoni* experiencing higher salinity and greater sea ice concentration and thickness, compared to post-2000 cohorts which were born during a period of decreasing salinity and sea ice (Figure 5, Figure S13). Historical regionally restricted sea ice losses, combined with recent strong anomalies and circumpolar decreases, may be increasing selection pressure on *D. mawsoni* to produce eggs that are able to float to the surface even in fresher, more stratified environments with reduced salinity due to sea ice loss. Evidence for such directional selection may be observed in the results of our functional enrichment analysis (Figure 4, Table S5, Table S6), where we identify multiple Gene Ontology (GO) terms across all categories that play a role in the osmoregulatory pathways implicated in determining egg buoyancy. Precedent exists for genomic adaptation to changing salinity regimes in fish, mainly relating to species with distributions between the more saline environments of the North Sea and the brackish waters of the Baltic Sea such as Atlantic cod (*Gadus morhua*) (Berg et al., 2015), Atlantic herring (*Clupea harengus*) (Guo et al., 2016; Andersson et al., 2023), and three-spined stickleback (*Gasterosteus aculeatus*) (Guo et al., 2015). With respect to freshening in response to sea ice loss, no studies yet exist investigating genomic adaptation to sea ice loss in vertebrates, though there is a growing body of literature revealing adaptive responses in polar microbes and diatoms to salinity changes linked to sea ice (Ewert and Deming, 2013; Dawson et al., 2020; Freyria et al., 2022; Sadler et al., 2025).

**Figure 5.**
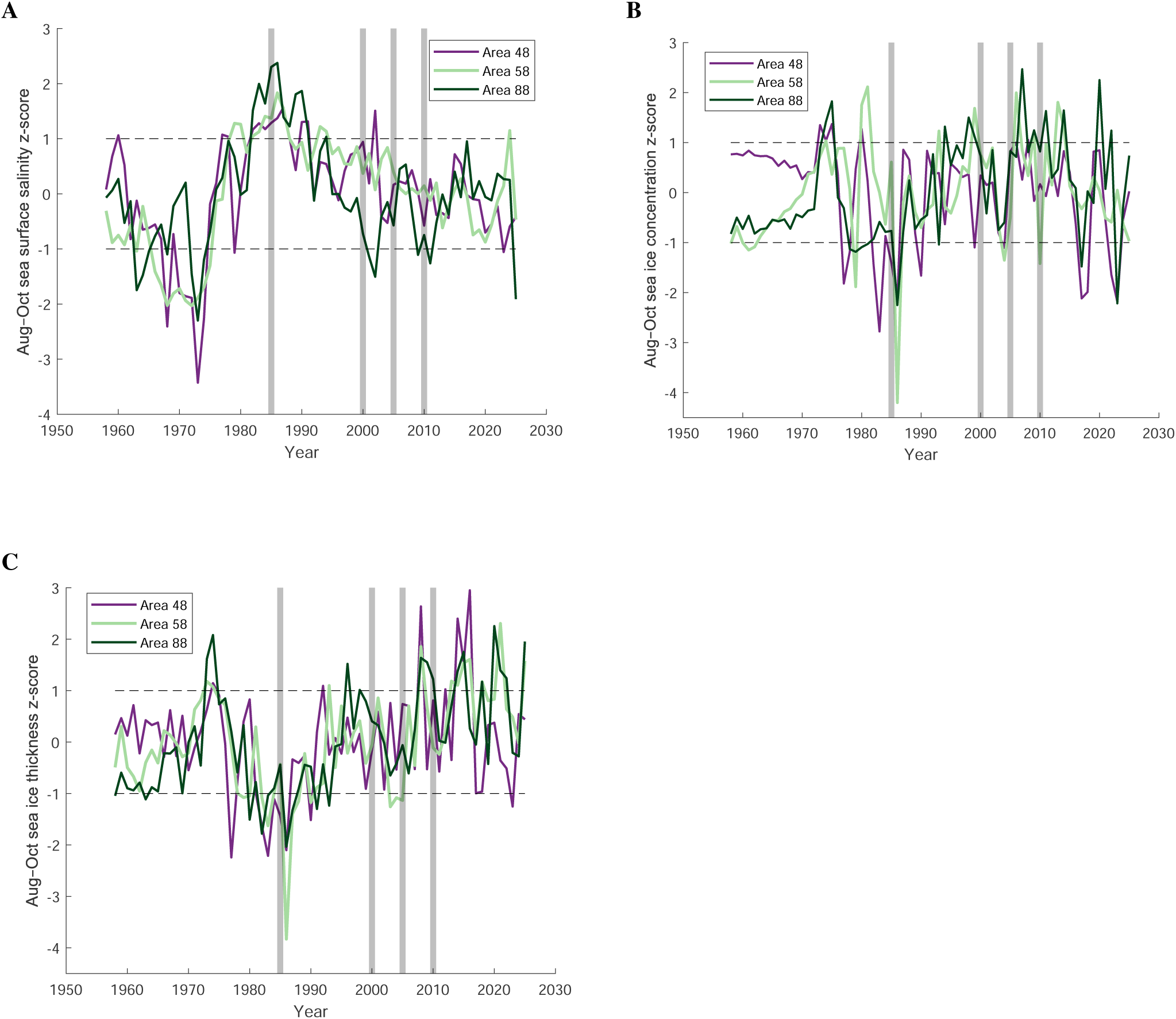
Time series based on z-scores of (A) sea surface salinity, (B) sea ice concentration, and (C) sea ice thickness. Z-scores were derived by taking anomaly values (calculated by subtracting from each point the time series mean over all years from 1958 to 2025), and dividing these by their standard deviation. Raw data from the ORAS5 global ocean reanalysis. Raw data averaged temporally over the estimated *D. mawsoni* egg incubation period (August–October), and spatially over the hypothesized *D. mawsoni* spawning grounds within each CCAMLR area (Figure 1). Dashed black lines correspond to one standard deviation from the mean. The purple line show data from CCAMLR area 48, the light green line from CCAMLR area 58, and the dark green line from CCAMLR area 88. Shaded gray bars highlight the estimated birth years for *D. mawsoni* used in this study.

### 4.3 Implications for fisheries management

Integration of outcomes from genomic analyses into management remains aspirational in most of the world’s fisheries (Bernos et al., 2020). While genomics has often been cited as a critical tool in distinguishing between stocks, assessing the connectivity between and ultimately health of regional populations, and identifying adaptive responses (Andersson et al., 2024), it is infrequently implemented into actual management practices (Bernatchez et al., 2017). The fishery for *D. mawsoni* is regulated by the Commission for the Conservation of Antarctic Marine Living Resources (CCAMLR), an international body composed of 27 member countries charged with management of Southern Ocean fisheries in the high seas to facilitate conservation via a precautionary, ecosystem-based approach (CCAMLR, 1980). In recent years, CCAMLR members have emphasized the need for greater integration of climate change into their management strategy (Cavanagh and Pardo, 2023). Our results highlighting the connection between genomic adaptation and salinity imply that *D. mawsoni* stocks experiencing greater reductions in salinity due to sea ice loss will be more vulnerable to added pressure from exploitation, and should thus be subject to more precautionary management measures.

In the Ross Sea region (CCAMLR management area 88), the largest area of *D. mawsoni* exploitation outside of national Exclusive Economic Zones, catch limits are determined based on the stock assessment model CASAL (Bull et al., 2012), which uses age or size input data and other population parameters (e.g., maturity, tag-release data, sex ratios) to create model outputs (Grüss et al., 2021). CCAMLR decision rules dictate that spawning biomass should not drop below certain pre-established pre-exploitation levels during the 35-year period over which the CASAL model projects (Delegation of the United Kingdom, 2019). Similar to biomass trend estimates from other regions of the Southern Ocean, the CASAL model does not consider climate change impacts or distribution shifts over the 35-year projection period (Delegation of the United Kingdom, 2019). A CASAL2 model has been in development in recent years, with the aim to increase flexibility in inputting population dynamics, allowing qualitative impacts such as population connectivity, adaptive pressure, and distribution shifts to be accounted for over the 35-year projection period (Dunn and Rasmussen, 2021). Conceivably, a salinity parameter could be added to the CASAL2 model to allow for modulation of catch limits during years when increased sea ice loss results in greater reductions in salinity, reducing water density, and thus reducing the probability that *D. mawsoni* eggs reach the surface, where they are more likely to survive and contribute to recruitment.

Our results establish an important precedent on which future studies based on samples for which greater certainty exists regarding age and early life locations may build. Indeed, now that we identified a connection between genomic adaptation and salinity using GEA, it would be of interest to investigate the extent to which the genomic makeup of existing *D. mawsoni* populations is maladapted to future salinity conditions, using a genomic offset analysis (Lind et al., 2024). Analyses of future maladaptation will be critical to any precautionary management strategy that requires updating stock assessment models, which often project over >30 years, from the capacity to only consider past and present impacts of climate change, to also including impacts based on future projections (Murphy et al., 2025).

## Data availability

Data files, commands, and scripts for various bioinformatics and statistical analyses (NOT included in Caccavo et al. 2025a) are be accessible from the following GitHub repository: https://github.com/jcaccavo/TOA_clim_gen.

## Author contributions

JAC conceived of and designed the research. MG provided expert advice on the selection of data to represent environmental exposure. JAC created the SNP dataset used from Caccavo et al. (2025). JAC performed the genotype-environment association (GEA) and gene ontology (GO) analyses with support from EC and PdV. JAC drafted the manuscript with input from all authors. All authors read and approved the manuscript.

## Acknowledgements

JAC acknowledges support from a postdoctoral fellowship adminstered by the Institut Pierre-Simon Laplace (IPSL), made possible by French state aid managed by the ANR (‘Agence Nationale de la Recherche’) under the ‘Investissements d’avenir’ program, with the reference ANR-11-IDEX-0004-17-EURE-0006. We thank Rebecca Konijnenberg and Christopher Jones for their input on the choice of fishing data type to use in the RDA. We also thank Steve Parker for his advice regarding the selection of hypothesized spawning grounds for *D. mawsoni*.

## Supplementary Tables

**Table S1.** Sample metadata including type, management area and subarea, longitude, latitude, year of capture, estimated year of birth, cohort, total length (cm), and final coverage. na, not available.

| Sample | Type | Area | Subarea | Longitude | Latitude | Year of capture | Estimated birth year | Cohort | Length (cm) | Coverage |
| --- | --- | --- | --- | --- | --- | --- | --- | --- | --- | --- |
| 9432 | tissue | 48 | 48.1 | -58.00 | -62.00 | 2001 | 1985 | pre-2000 | na | 11.40 |
| 312 | otolith | 48 | 48.5 | -27.84 | -74.64 | 2013 | 2000 | post-2000 | 123.0 | 12.46 |
| 313 | otolith | 48 | 48.5 | -27.84 | -74.64 | 2013 | 2000 | post-2000 | 118.0 | 5.92 |
| 314 | otolith | 48 | 48.5 | -27.84 | -74.64 | 2013 | 2000 | post-2000 | 103.0 | 13.27 |
| 315 | otolith | 48 | 48.5 | -27.84 | -74.64 | 2013 | 2000 | post-2000 | 128.0 | 10.86 |
| 527 | otolith | 48 | 48.2 | -38.72 | -61.56 | 2018 | 2005 | post-2000 | 151.0 | 6.76 |
| 168 | otolith | 48 | 48.2 | -34.74 | -61.40 | 2019 | 2005 | post-2000 | 158.0 | 6.92 |
| 9479 | tissue | 58 | 58.4.2 | 44.00 | -67.20 | 2001 | 1985 | pre-2000 | na | 6.43 |
| 412 | otolith | 58 | 58.4.2 | 71.74 | -66.49 | 2019 | 2010 | post-2000 | 105.0 | 6.88 |
| 419 | otolith | 58 | 58.4.2 | 71.48 | -66.60 | 2019 | 2010 | post-2000 | 89.5 | 12.03 |
| 433 | otolith | 58 | 58.4.2 | 71.60 | -66.44 | 2019 | 2005 | post-2000 | 122.5 | 11.07 |
| 9506 | tissue | 88 | 88.1 | 179.00 | -77.00 | 2001 | 1985 | pre-2000 | na | 7.29 |
| 9512 | tissue | 88 | 88.1 | 179.00 | -77.00 | 2001 | 1985 | pre-2000 | na | 22.13 |
| 9522 | tissue | 88 | 88.1 | 171.00 | -73.00 | 2001 | 1985 | pre-2000 | na | 9.52 |
| 9525 | tissue | 88 | 88.1 | 176.00 | -71.00 | 2001 | 1985 | pre-2000 | na | 7.94 |
| 9540 | tissue | 88 | 88.1 | 179.00 | -70.00 | 2001 | 1985 | pre-2000 | na | 13.40 |
| 9544 | tissue | 88 | 88.1 | 179.00 | -70.00 | 2001 | 1985 | pre-2000 | na | 5.90 |
| 1-425 | tissue | 88 | 88.1 | 178.00 | -64.00 | 2019 | 2005 | post-2000 | 148.0 | 15.37 |
| 33-429 | tissue | 88 | 88.1 | 181.00 | -72.00 | 2019 | 2005 | post-2000 | 156.0 | 13.09 |
| 34-430 | tissue | 88 | 88.1 | 181.00 | -72.00 | 2019 | 2005 | post-2000 | 142.0 | 11.82 |
| 35-431 | tissue | 88 | 88.1 | 181.00 | -72.00 | 2019 | 2005 | post-2000 | 152.0 | 8.78 |
| 4-456 | tissue | 88 | 88.1 | 176.00 | -65.00 | 2019 | 2005 | post-2000 | 145.0 | 19.92 |
| 42-438 | tissue | 88 | 88.1 | 180.00 | -73.00 | 2020 | 2005 | post-2000 | 127.0 | 11.28 |
| 50-446 | tissue | 88 | 88.1 | 180.00 | -73.00 | 2020 | 2005 | post-2000 | 158.0 | 27.09 |

**Table S2.** Environmental variables, units, and descriptions from the ORAS5 global ocean reanalysis used in the genotype–environment association (GEA) analysis. Monthly averages for August, September, and October (*D. mawsoni* spawning months) were downloaded as NetCDF4 files from the Copernicus Marine Service (CMEMS) for respective birth (1985, 2000, 2005, and 2010) and catch (2001, 2013, 2018, 2019, and 2020) years depending on the environmental exposure scenario. All variables have an approximate horizontal resolution of 0.25° × 0.25°.

| Environmental variable | Units | Description |
| --- | --- | --- |
| Sea ice thickness | m | Mean thickness of the sea ice layer in the area of the grid cell covered by ice. |
| Sea ice concentration | dimensionless | Fraction of the area of the grid cell containing sea ice. |
| Mixed layer depth 0.01 | m | The depth of the ocean where the average sea water density exceeds the near-surface density plus $0.01 \text{ kg m}^{-3}$. |
| Mixed layer depth 0.03 | m | The depth of the ocean where the average sea water density exceeds the near-surface density plus $0.03 \text{ kg m}^{-3}$. |
| Sea surface salinity | PSU | Salt concentration close to the ocean surface. |
| Sea surface temperature | $^\circ\text{C}$ | Water temperature close to the ocean surface. |
| Meridional velocity | $\text{m s}^{-1}$ | Horizontal velocity of a water parcel as a vector quantity decomposed into the meridional (along the y-axis) component of the ocean model. |
| Zonal velocity | $\text{m s}^{-1}$ | Horizontal velocity of a water parcel as a vector quantity decomposed into the zonal (along the x-axis) component of the ocean model. |

**Table S3.** Coordinates of bounding boxes defining hypothesized spawning grounds. Place names refer to locations defined on the map in Figure 1. Note that for spawning ground number 7 (Ross sea), the bounding box was split into two sets of coordinates because they occur on either side of the 180 meridian. min, minimum; max, maximum; lon, longitude; lat, latitude.

| CCAMLR area | Spawning ground number | Place names | Min lon | Max lon | Min lat | Max lat |
| --- | --- | --- | --- | --- | --- | --- |
| 88 | 1 | Amundsen sea | -132.8 | -118.2 | -70.8 | -68.5 |
| 48 | 2 | Scotia sea | -27.8 | -24.5 | -60.7 | -58.1 |
| 58 | 3 | Cosmonaut sea | 31.8 | 35.2 | -66.8 | -64.5 |
| 58 | 4 | Banzare bank | 76.6 | 84.1 | -62.2 | -54.5 |
| 58 | 5 | Kerguelen plateau | 61.8 | 72.2 | -50.7 | -46.4 |
| 58 | 6 | Davis sea | 92.5 | 104.7 | -64.4 | -62.7 |
| 88 | 7 | Ross sea | 167.0 | 180.0 | -64.8 | -62.6 |
|  |  |  | -180.0 | 167.0 | -65.8 | -62.9 |

**Table S4.**
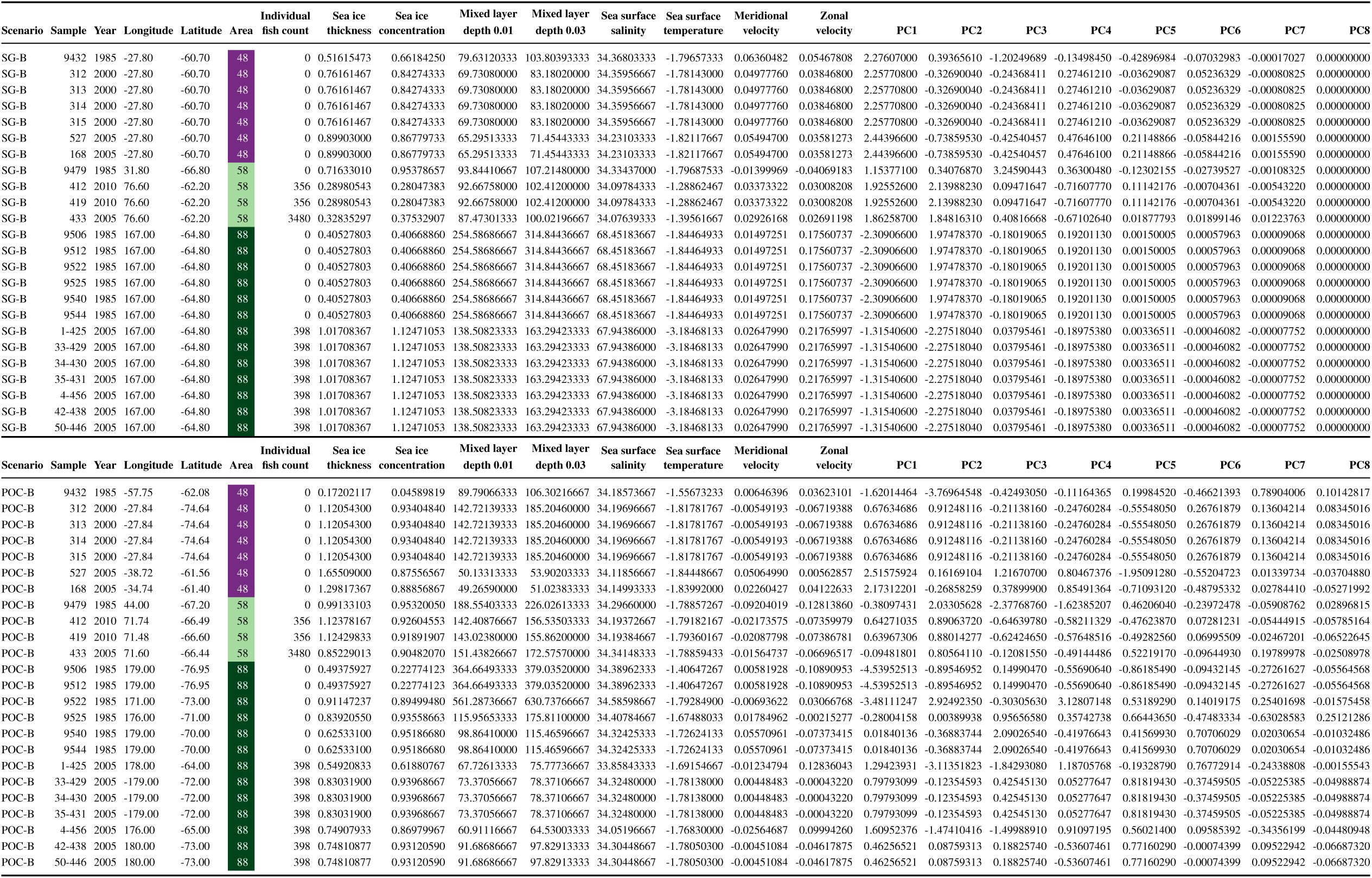
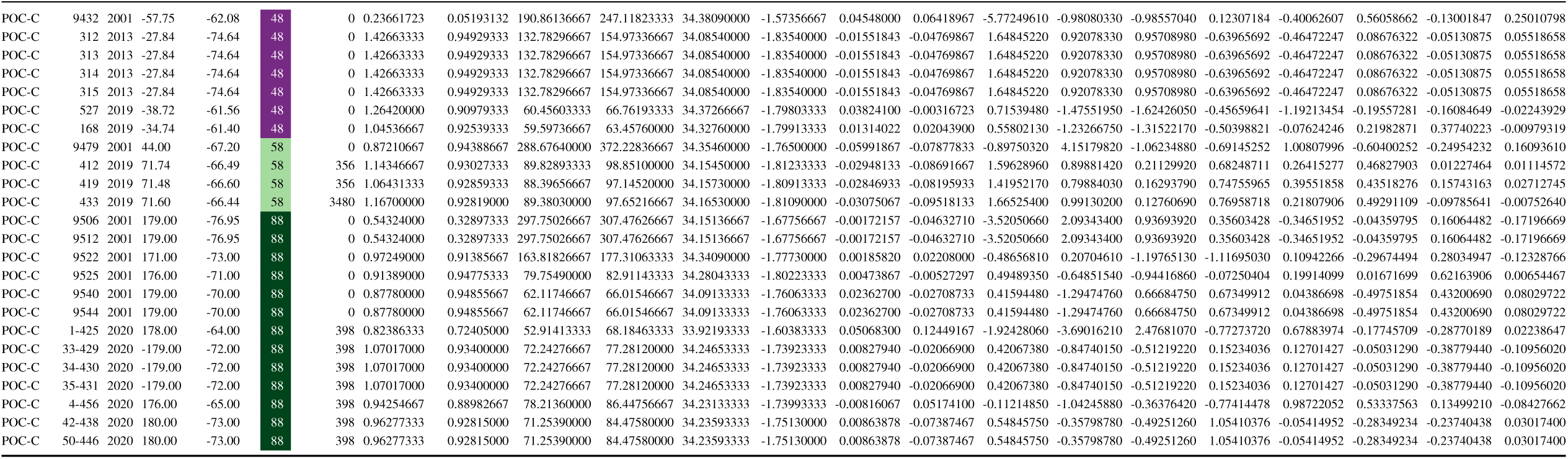
Input data used for pairs plots and principal component analysis (PCA) (raw environmental data, columns “Sea ice thickness” through “Zonal velocity”), and input data used for genotype-environment association (GEA) analysis (principal component (PC) scores, columns “PC1” through “PC3”). PC scores shown in columns “PC4” through “PC8” were generated by the PCA, but not used in the GEA. Scenario refers to Point of Capture–Time of Capture (POC-C), Spawning Ground–Time of Birth (SG-B), Point of Capture–Time of Birth (POC-B). Year refers to the year for which data were extracted: estimated birth year for the SG-B and POC-B scenarios; year of capture for the POC-C scenario. Area refers to fisheries management areas (48, 58, or 88). Individual fish count refers to the number of fish caught during the same year and over the same area from which environmental data were extracted; these data were used as the conditioning variable in the redundancy analysis (RDA).

**Table S5.** Gene ontology (GO) terms significantly (*p* > 0.01) overrepresented with respect to the whole genome. GO terms are derived from high-confidence GEA loci associated to predictor variable PC1 (largely driven by salinity, Figure S11a) from the Spawning Ground–Time of Birth (SG-B) scenario.

| GO category | GO ID | GO term | p-value | Fold enrichment | FDR |
| --- | --- | --- | --- | --- | --- |
| Molecular function | GO:0005509 | calcium ion binding | 0.000325894 | 1.636669150 | 0.265929347 |
| Molecular function | GO:0000166 | nucleotide binding | 0.001102533 | 1.340280927 | 0.339566697 |
| Molecular function | GO:0005216 | monoatomic ion channel activity | 0.001248407 | 2.232440760 | 0.339566697 |
| Molecular function | GO:0005085 | guanyl-nucleotide exchange factor activity | 0.001925501 | 2.061516257 | 0.392802133 |
| Molecular function | GO:0016787 | hydrolase activity | 0.002466923 | 1.360472685 | 0.402601891 |
| Molecular function | GO:0015179 | L-amino acid transmembrane transporter activity | 0.004040543 | 5.485425868 | 0.549513789 |
| Molecular function | GO:0005245 | voltage-gated calcium channel activity | 0.006441844 | 3.585245665 | 0.699104971 |
| Molecular function | GO:0005262 | calcium channel activity | 0.006853970 | 2.732775568 | 0.699104971 |
| Molecular function | GO:0016301 | kinase activity | 0.008777117 | 1.393654946 | 0.792429306 |
| Cellular component | GO:0034702 | monoatomic ion channel complex | 0.002774832 | 2.404346149 | 0.584403356 |
| Cellular component | GO:0005576 | extracellular region | 0.003730098 | 1.404395380 | 0.584403356 |
| Cellular component | GO:0005794 | Golgi apparatus | 0.005540516 | 1.512734452 | 0.584403356 |
| Cellular component | GO:0005737 | cytoplasm | 0.007517577 | 1.141924893 | 0.584403356 |
| Cellular component | GO:0000145 | exocyst | 0.008523937 | 5.971320206 | 0.584403356 |
| Biological process | GO:0034220 | monoatomic ion transmembrane transport | 0.002189739 | 1.831688953 | 1.000000000 |
| Biological process | GO:0007155 | cell adhesion | 0.002591516 | 1.714221944 | 1.000000000 |
| Biological process | GO:0003333 | amino acid transmembrane transport | 0.002928334 | 5.877332381 | 1.000000000 |
| Biological process | GO:0048513 | animal organ development | 0.003342516 | 2.279798572 | 1.000000000 |
| Biological process | GO:0060041 | retina development in camera-type eye | 0.004295296 | 2.379905718 | 1.000000000 |
| Biological process | GO:0006816 | calcium ion transport | 0.005139063 | 2.676804847 | 1.000000000 |
| Biological process | GO:0060287 | epithelial cilium movement involved in determination of left/right asymmetry | 0.007142623 | 6.258270591 | 1.000000000 |
| Biological process | GO:0006811 | monoatomic ion transport | 0.008303490 | 1.514000821 | 1.000000000 |
| Biological process | GO:0070588 | calcium ion transmembrane transport | 0.009645915 | 2.457793541 | 1.000000000 |

**Table S6.**
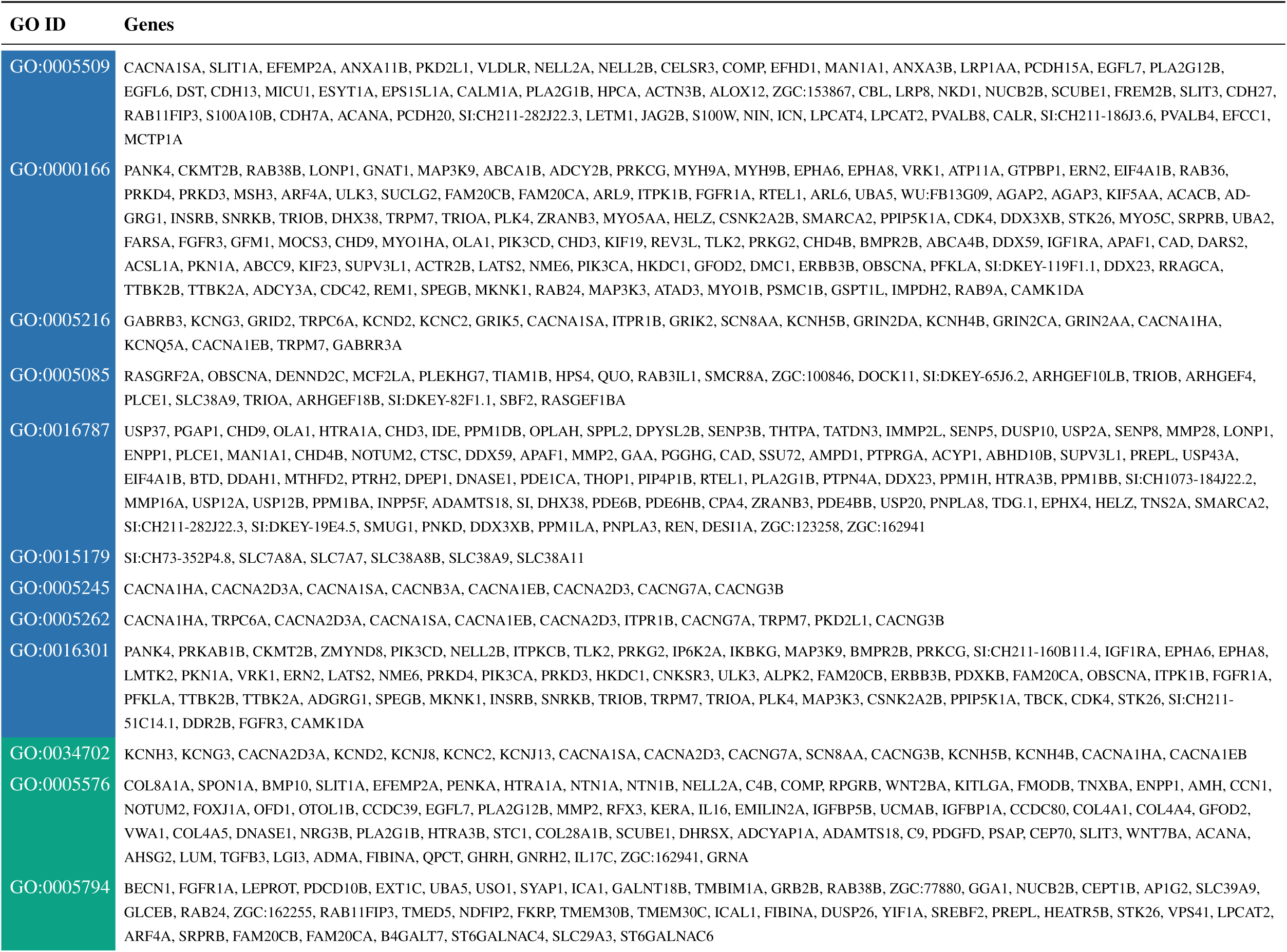
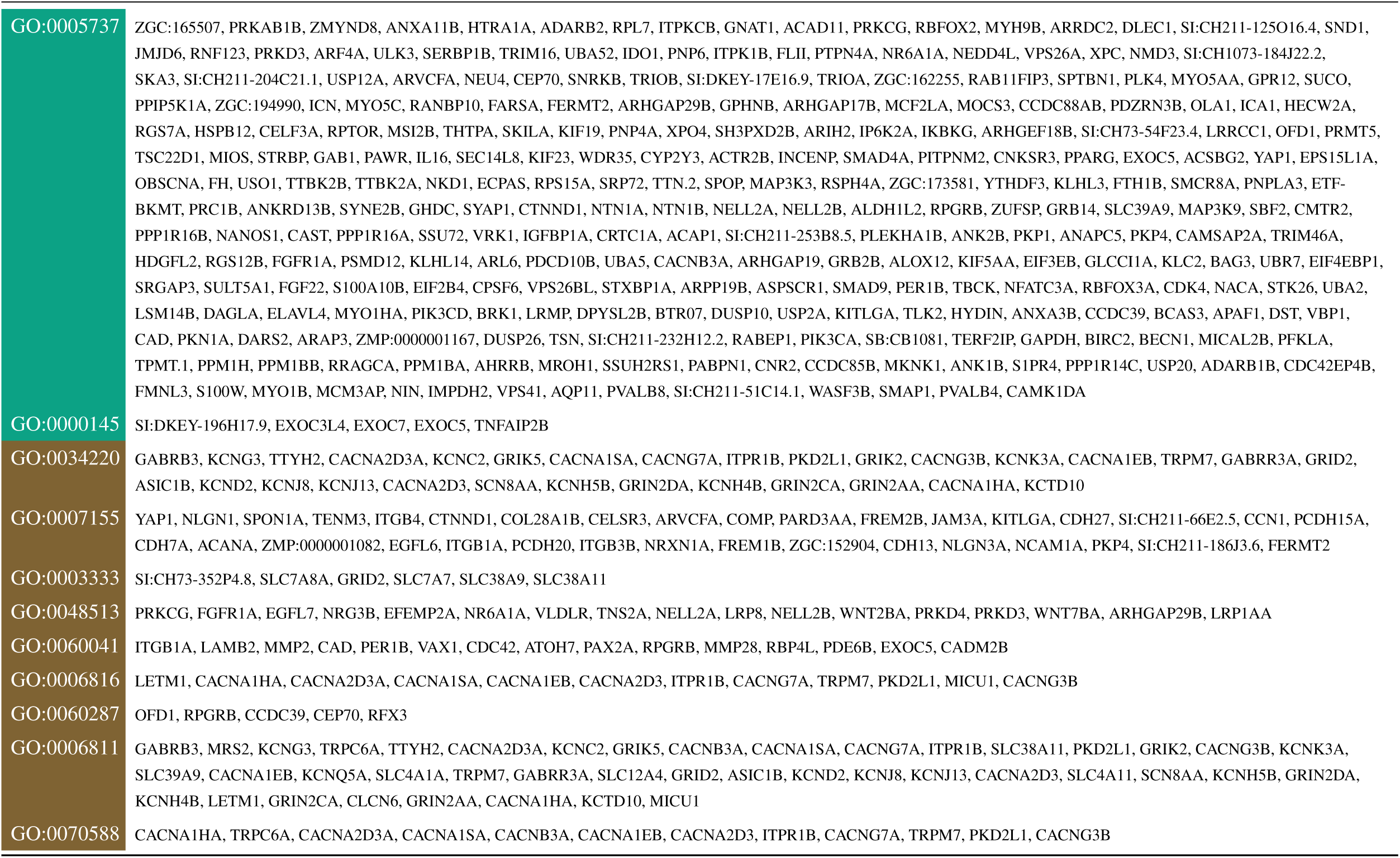
Genes associated with gene ontology (GO) terms, with GO IDs colour-coded by category: molecular function (blue), cellular component (green), or biological process (brown).

## Supplementary Figures

**Figure S1.**
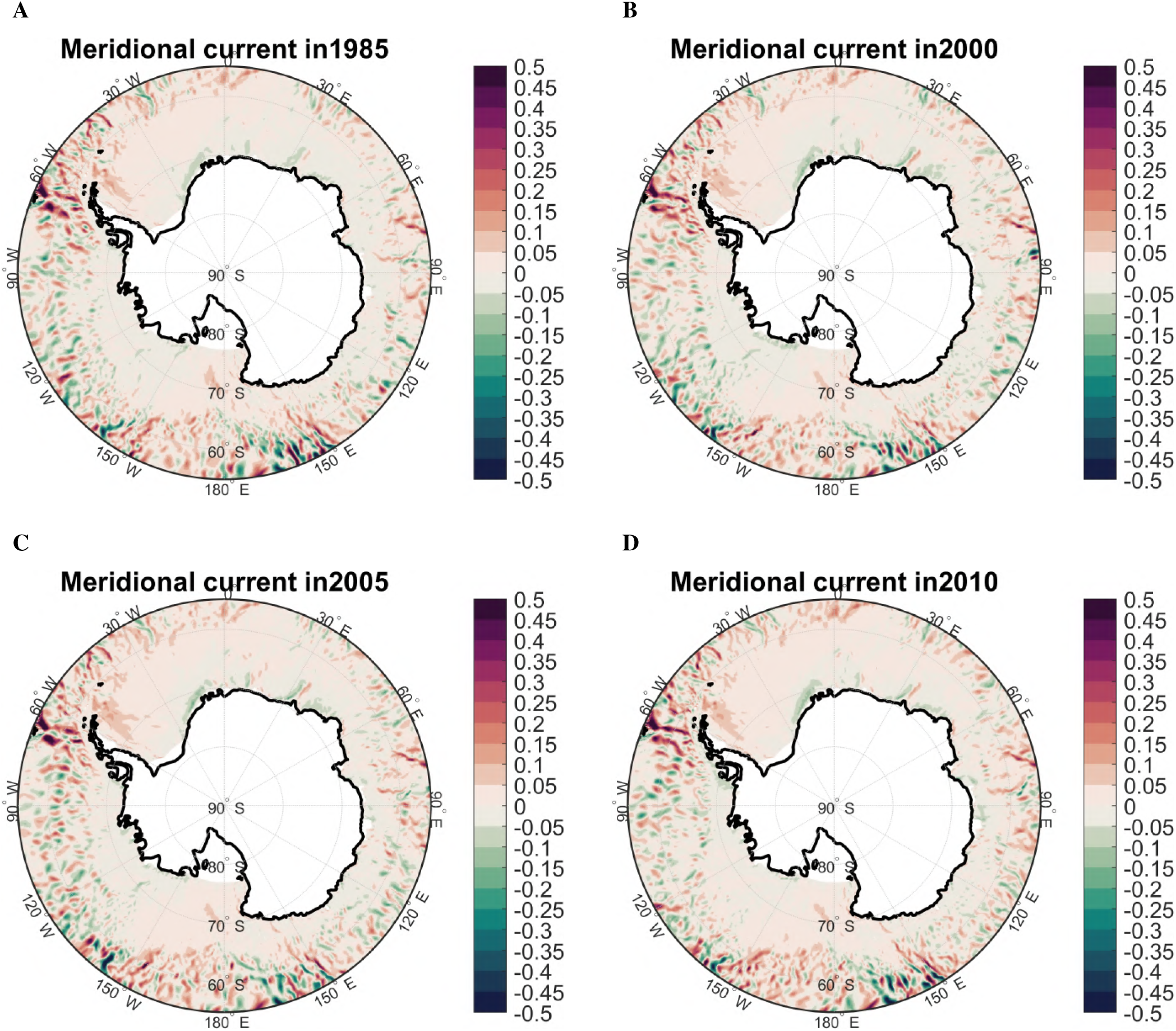
Map of meridional current velocity (m/s) across the Southern Ocean in (A) 1985, (B) 2000, (C) 2005, and (D) 2010.

**Figure S2.**
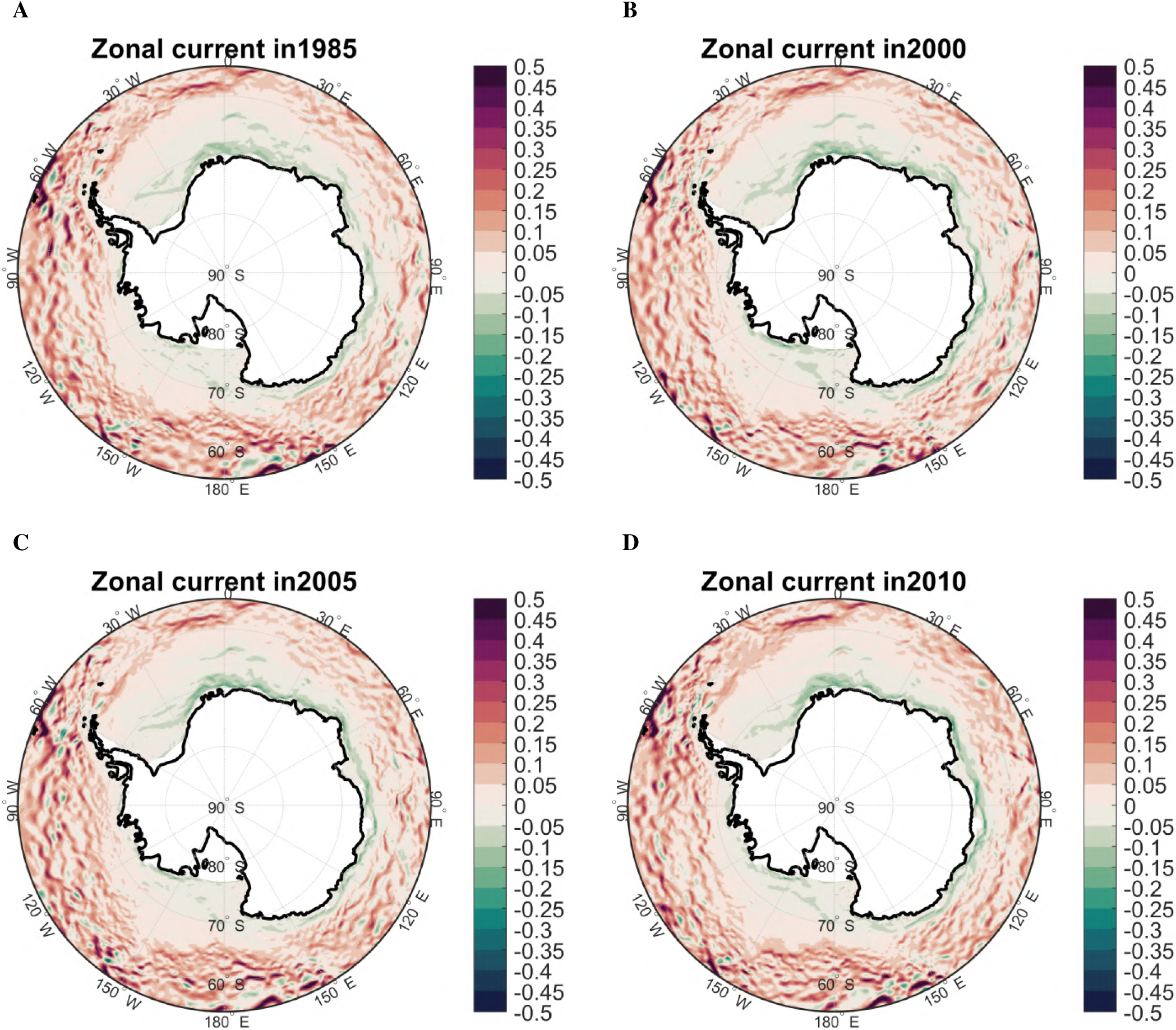
Map of zonal current velocity (m/s) across the Southern Ocean in (A) 1985, (B) 2000, (C) 2005, and (D) 2010.

**Figure S3.**
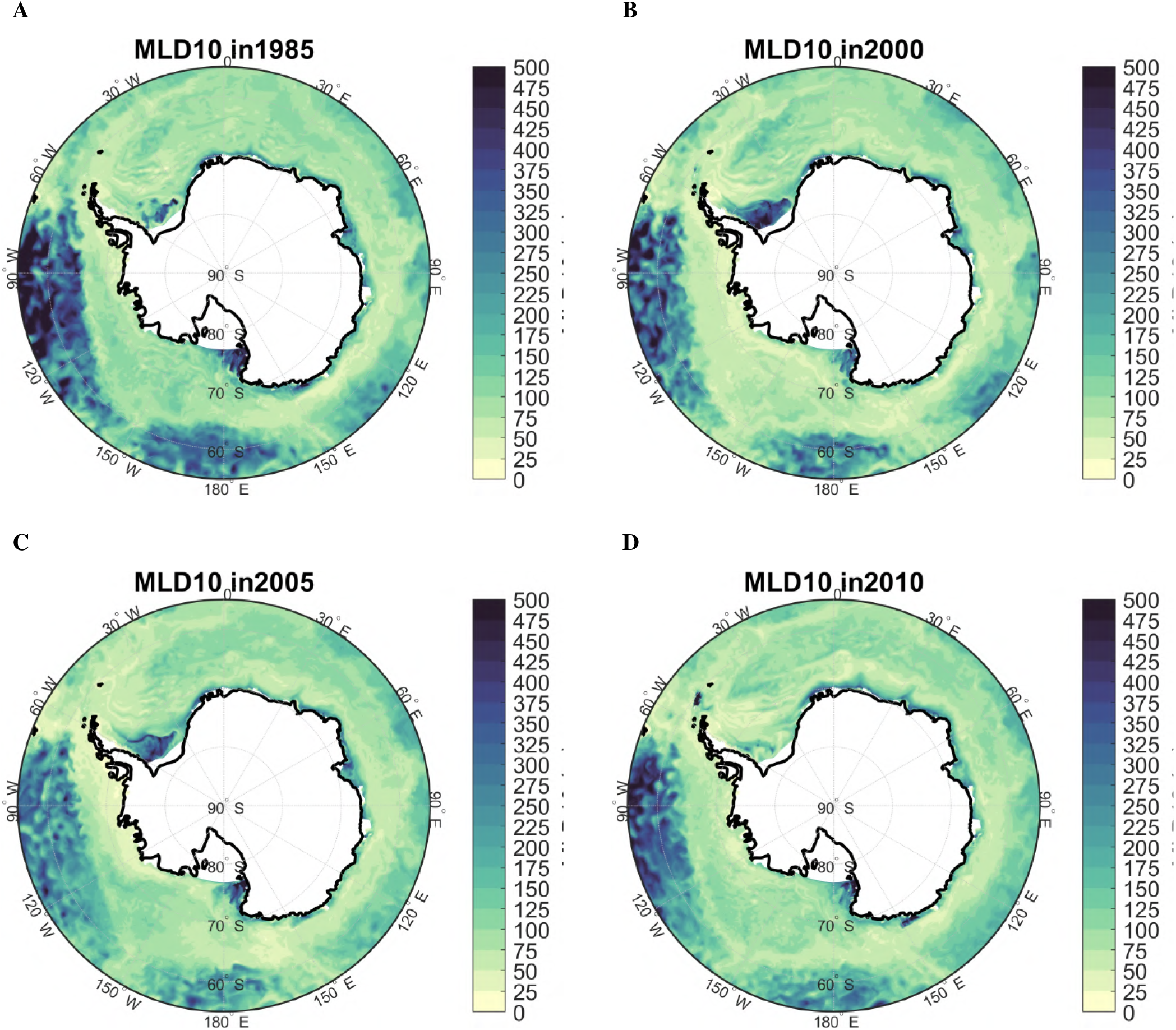
Map of mixed layer depth (m) at 0.01 density across the Southern Ocean in (A) 1985, (B) 2000, (C) 2005, and (D) 2010.

**Figure S4.**
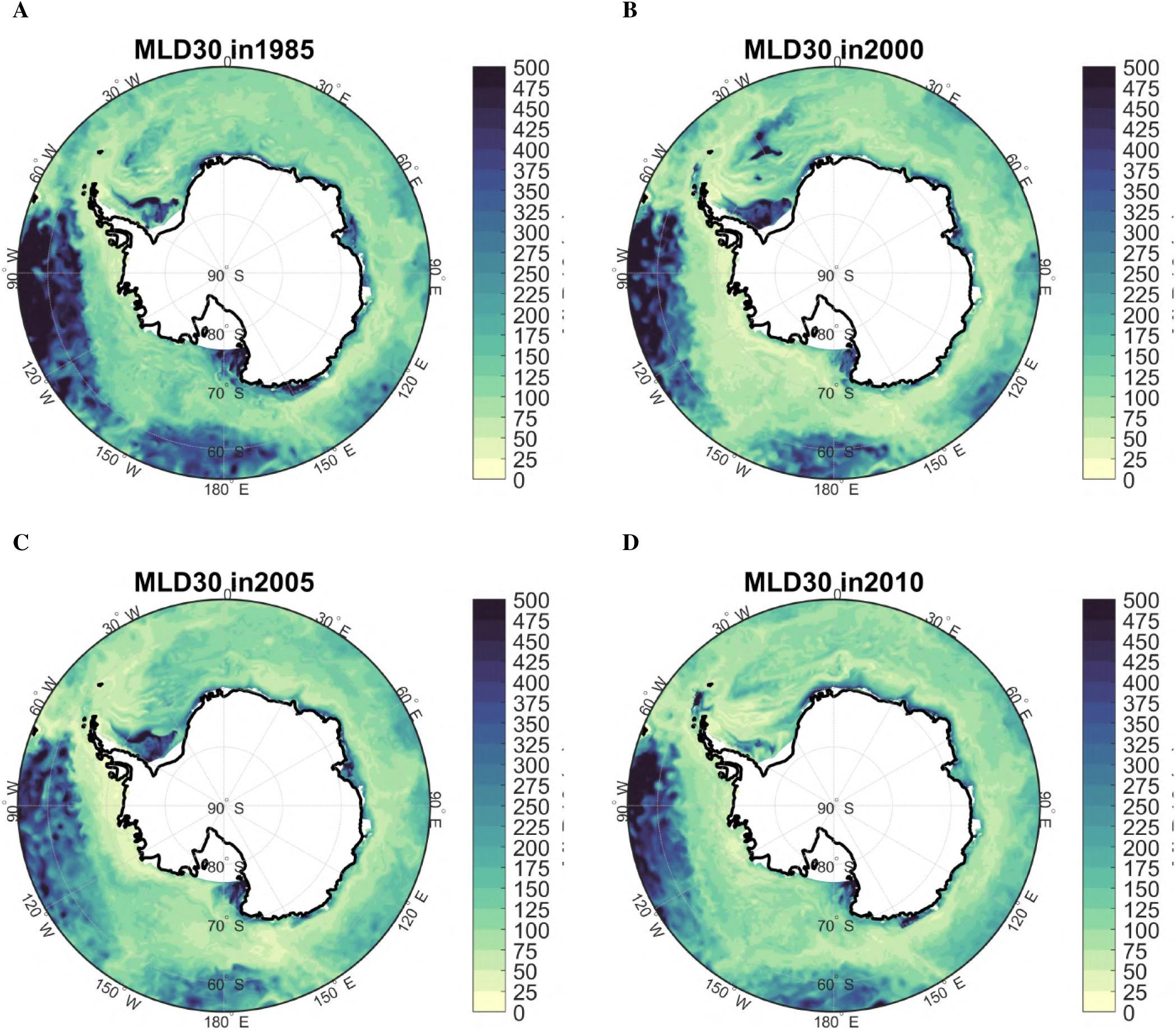
Map of mixed layer depth (m) at 0.03 density across the Southern Ocean in (A) 1985, (B) 2000, (C) 2005, and (D) 2010.

**Figure S5.**
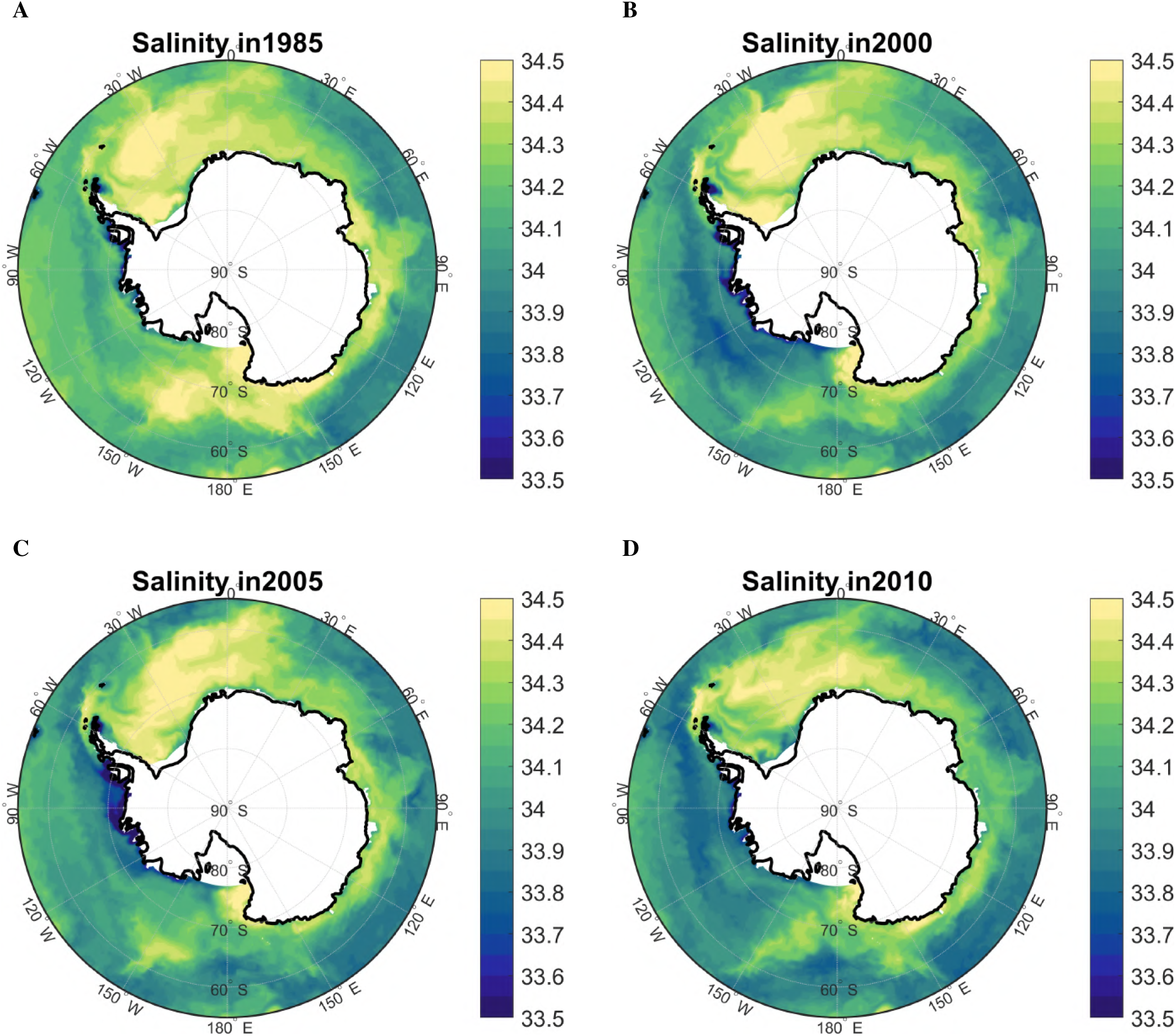
Map of salinty (PSU) across the Southern Ocean in (A) 1985, (B) 2000, (C) 2005, and (D) 2010.

**Figure S6.**
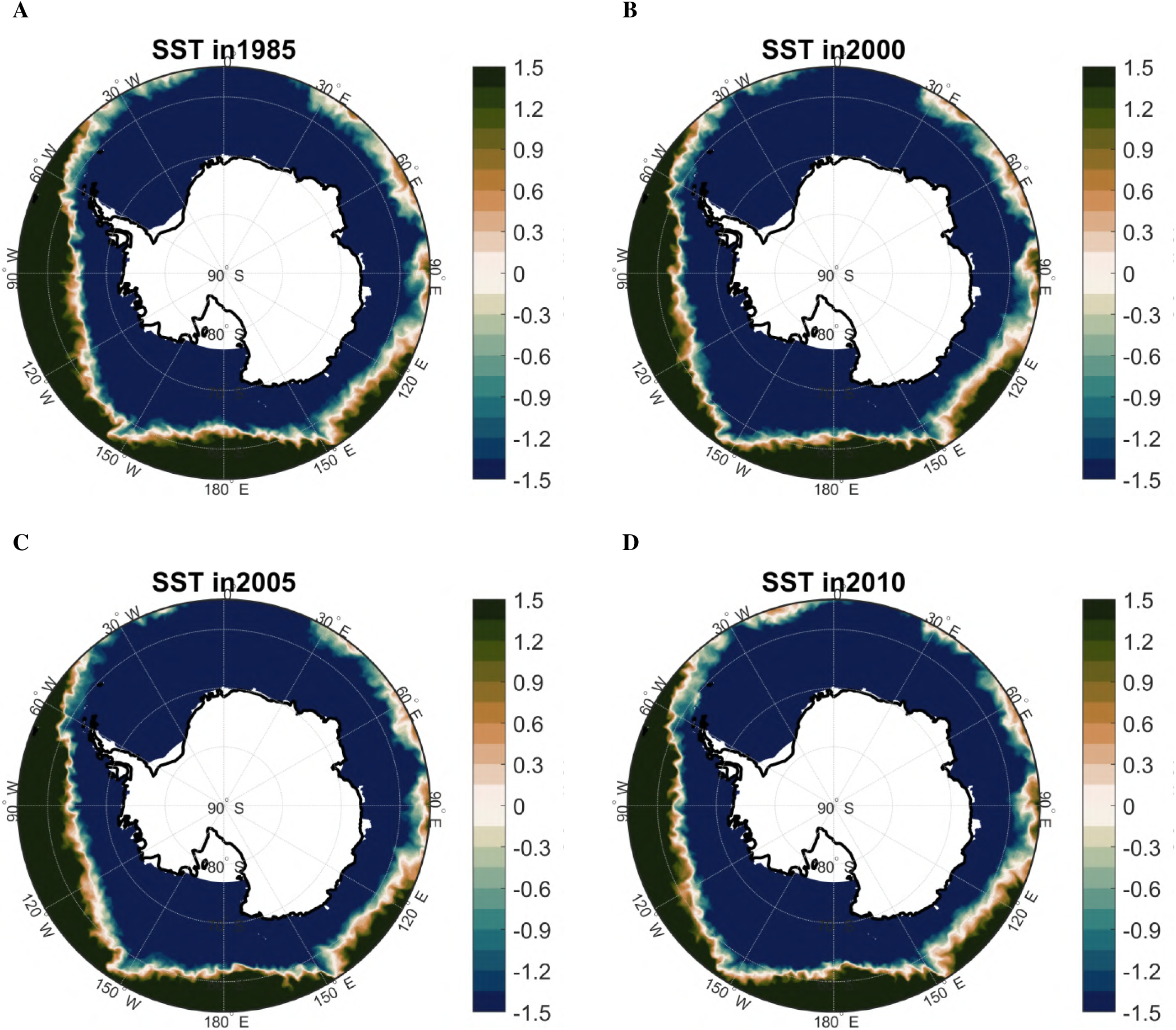
Map of sea surface temperature (°C) across the Southern Ocean in (A) 1985, (B) 2000, (C) 2005, and (D) 2010.

**Figure S7.**
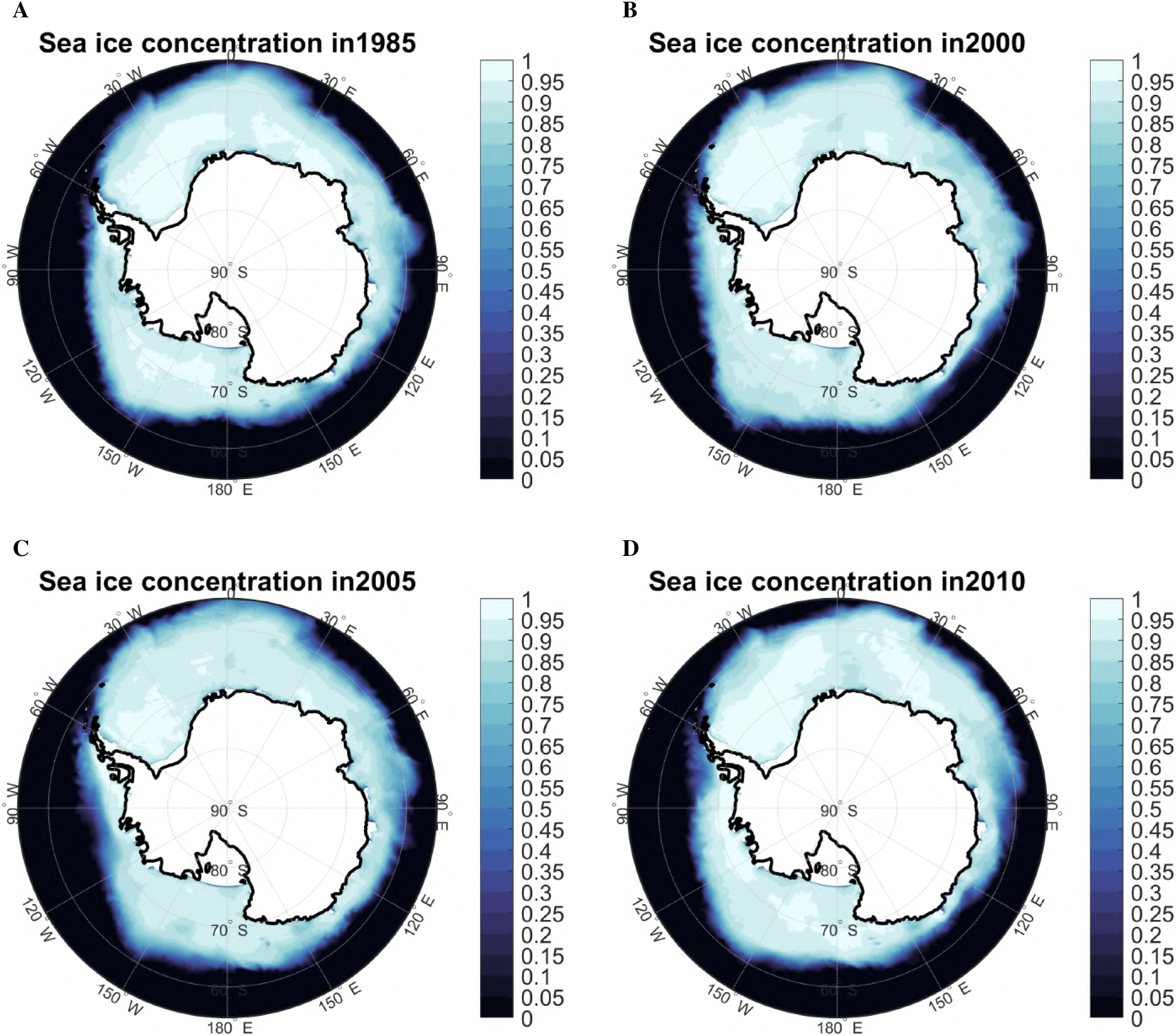
Map of sea ice concentration across the Southern Ocean in (A) 1985, (B) 2000, (C) 2005, and (D) 2010.

**Figure S8.**
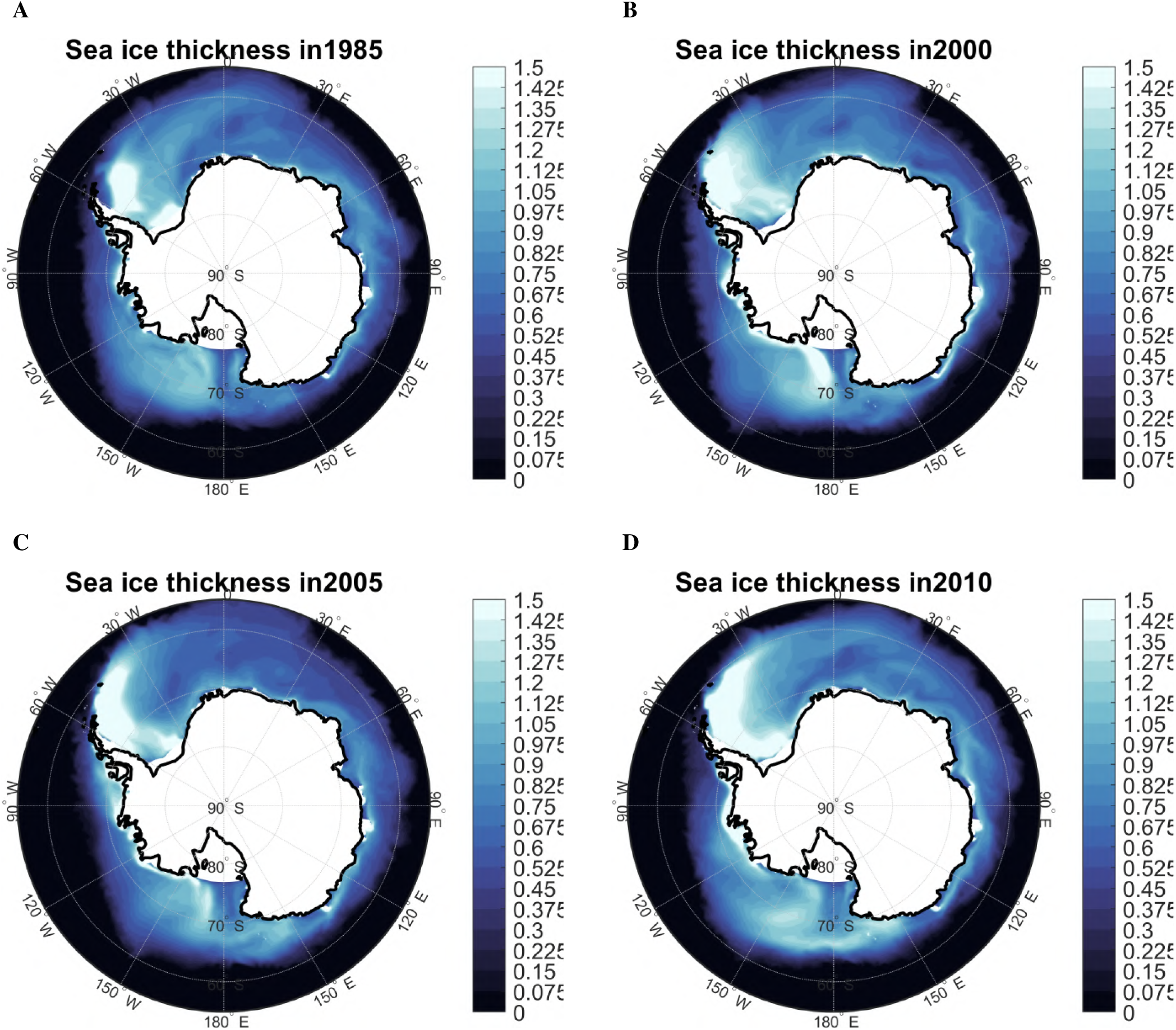
Map of sea ice thickness (m) across the Southern Ocean in (A) 1985, (B) 2000, (C) 2005, and (D) 2010.

**Figure S9.**
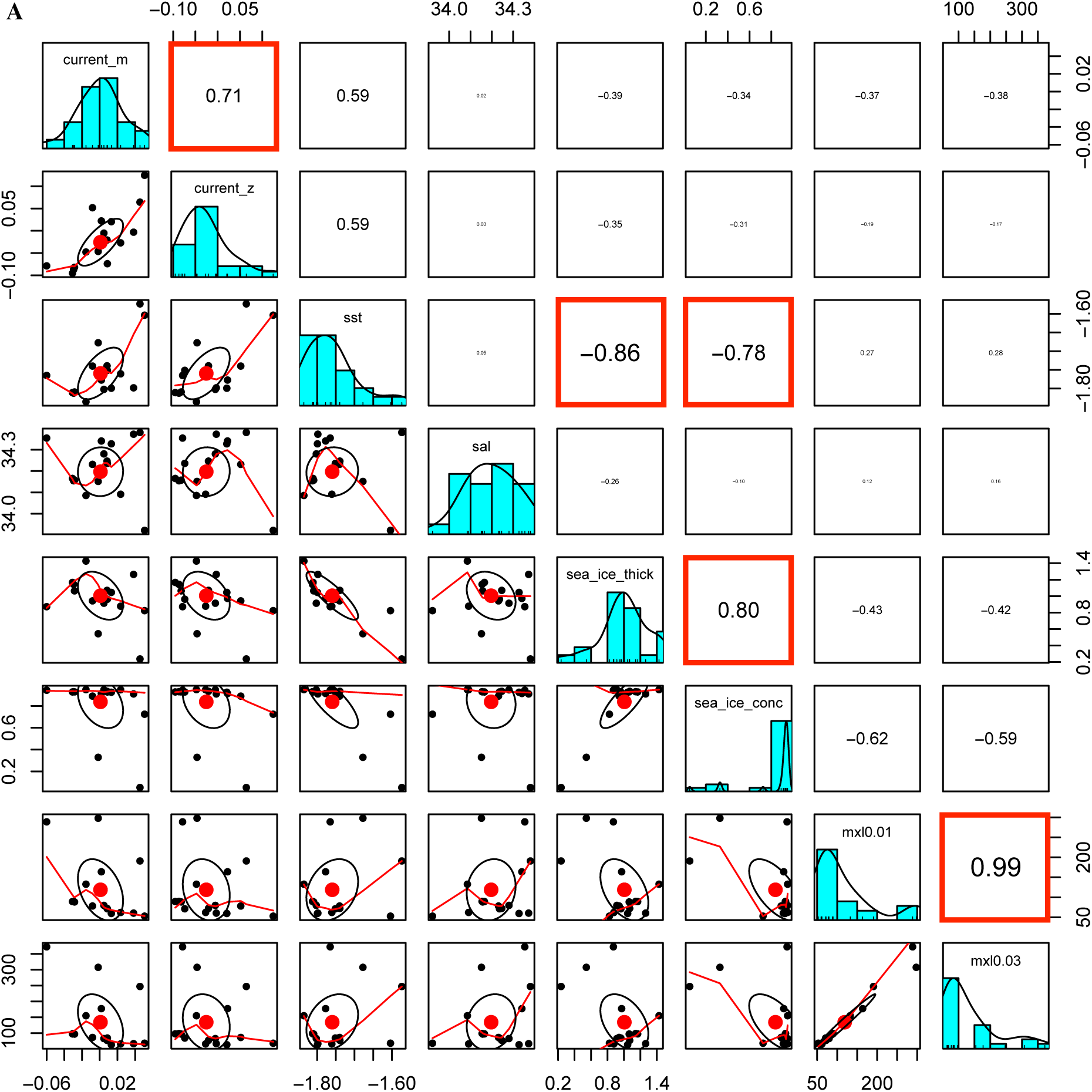
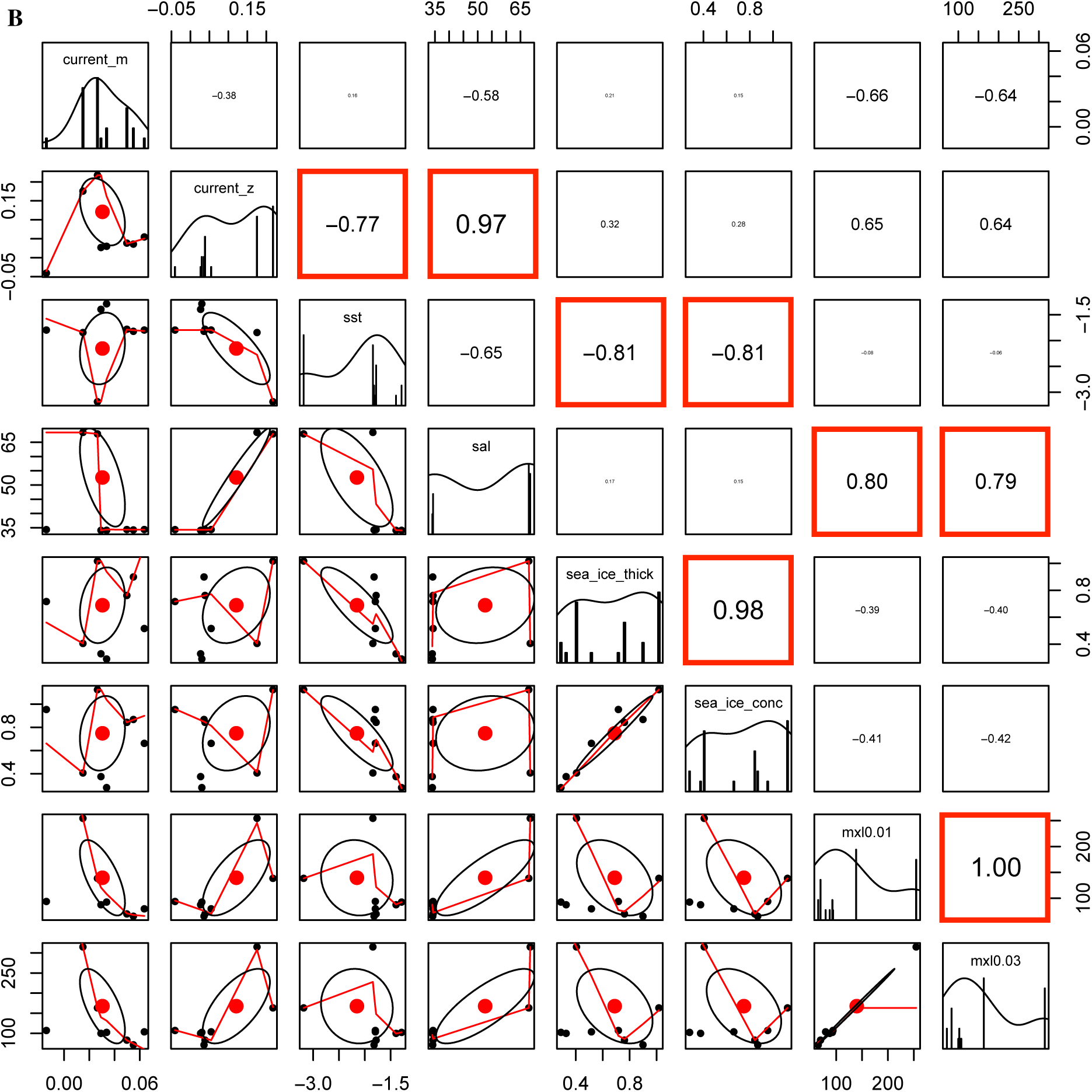
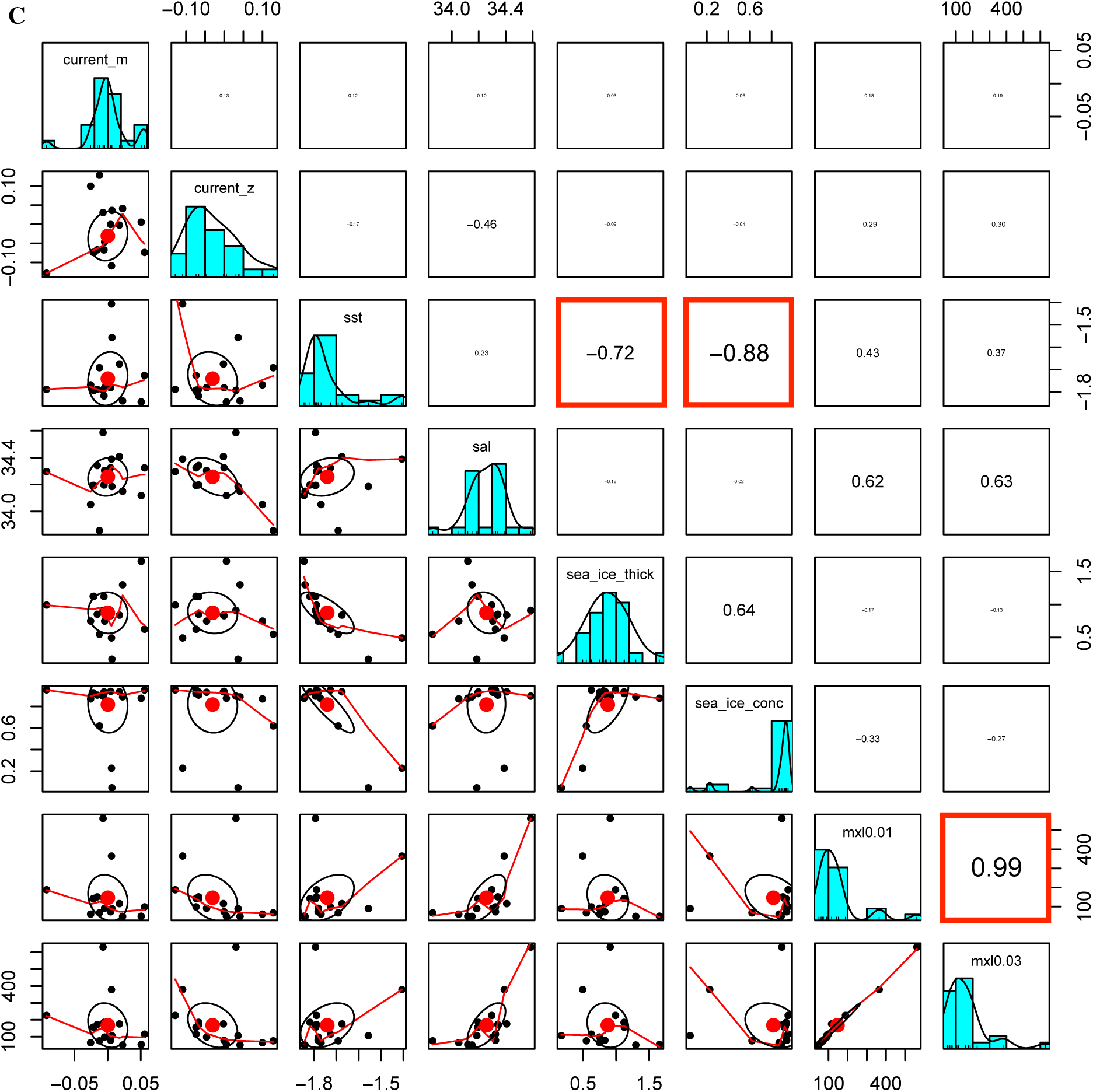
Pairs plots showing collinearity among environmental variables in scenarios (A) Point of Capture–Time of Capture (POC-C), (B) Spawning Ground–Time of Birth (SG-B), and (C) Point of Capture–Time of Birth (POC-B). *sea*_*ice*_*thick*, sea ice thickness; *sea*_*ice*_*conc*, sea ice concentration; *mxl*0.01, mixed-layer depth at 0.01 density; *mxl*0.03, mixed-layer depth at 0.03 density; *sal*, salinity; *sst*, sea surface temperature; *current*_*m*, meridional current velocity; *current*_*z*, zonal current velocity. Boxes with bold red outlines indicate significant collinearity (*r ≥ |*0.7|).

**Figure S10.**
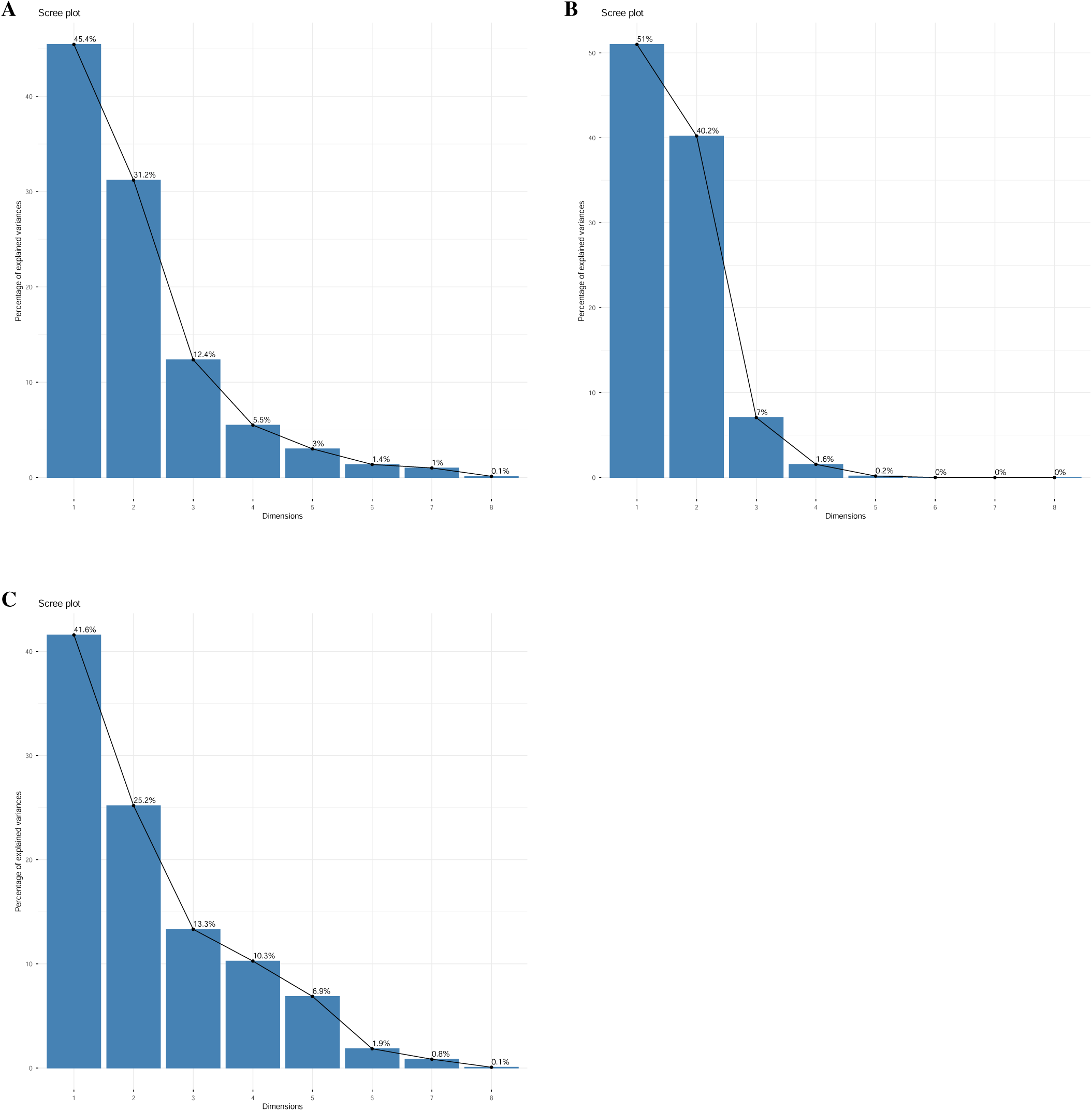
Scree plots from principal component analyses (PCAs) of all eight environmental variables from scenarios (A) Point of Capture– Time of Capture (POC-C), (B) Spawning Ground–Time of Birth (SG-B), and (C) Point of Capture–Time of Birth (POC-B).

**Figure S11.**
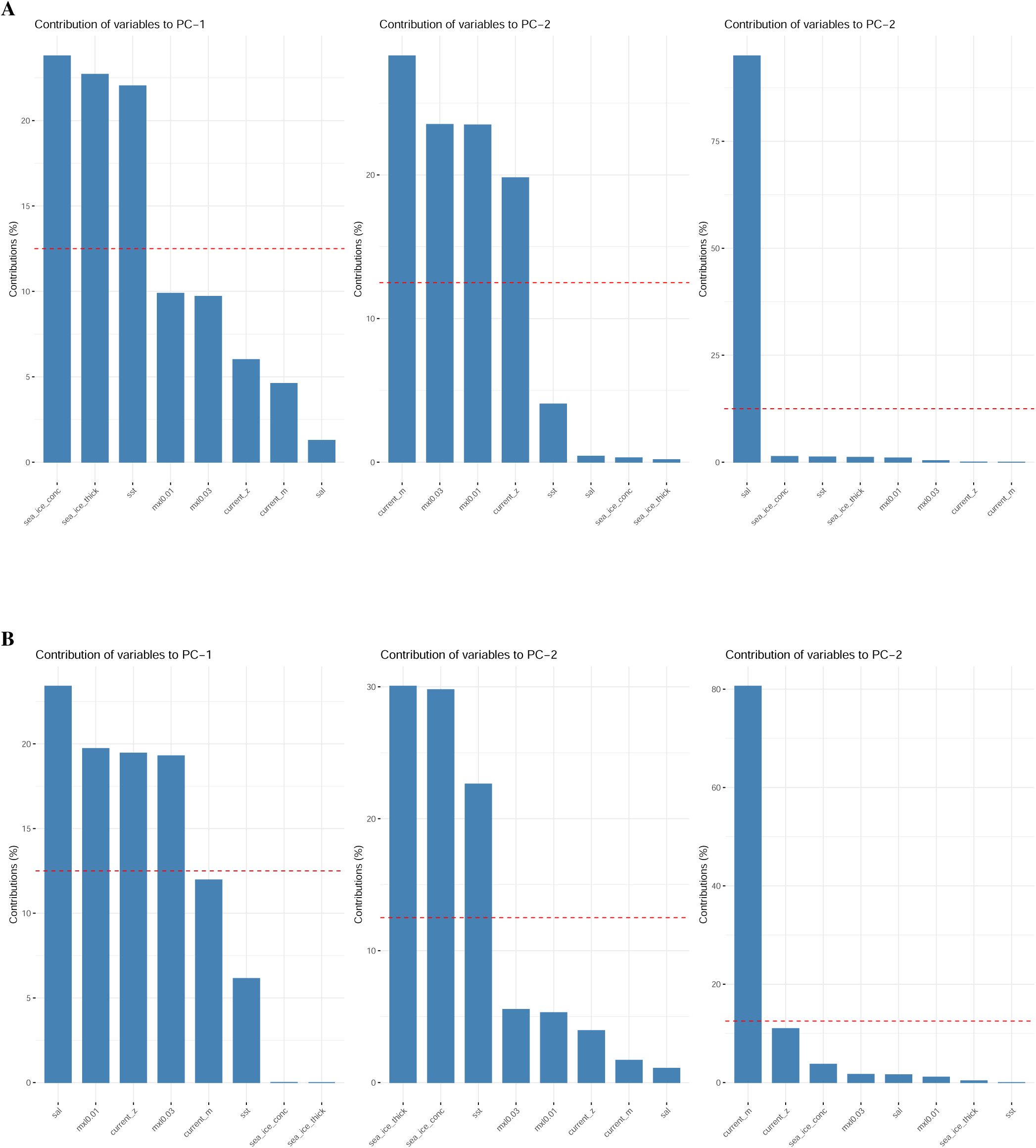
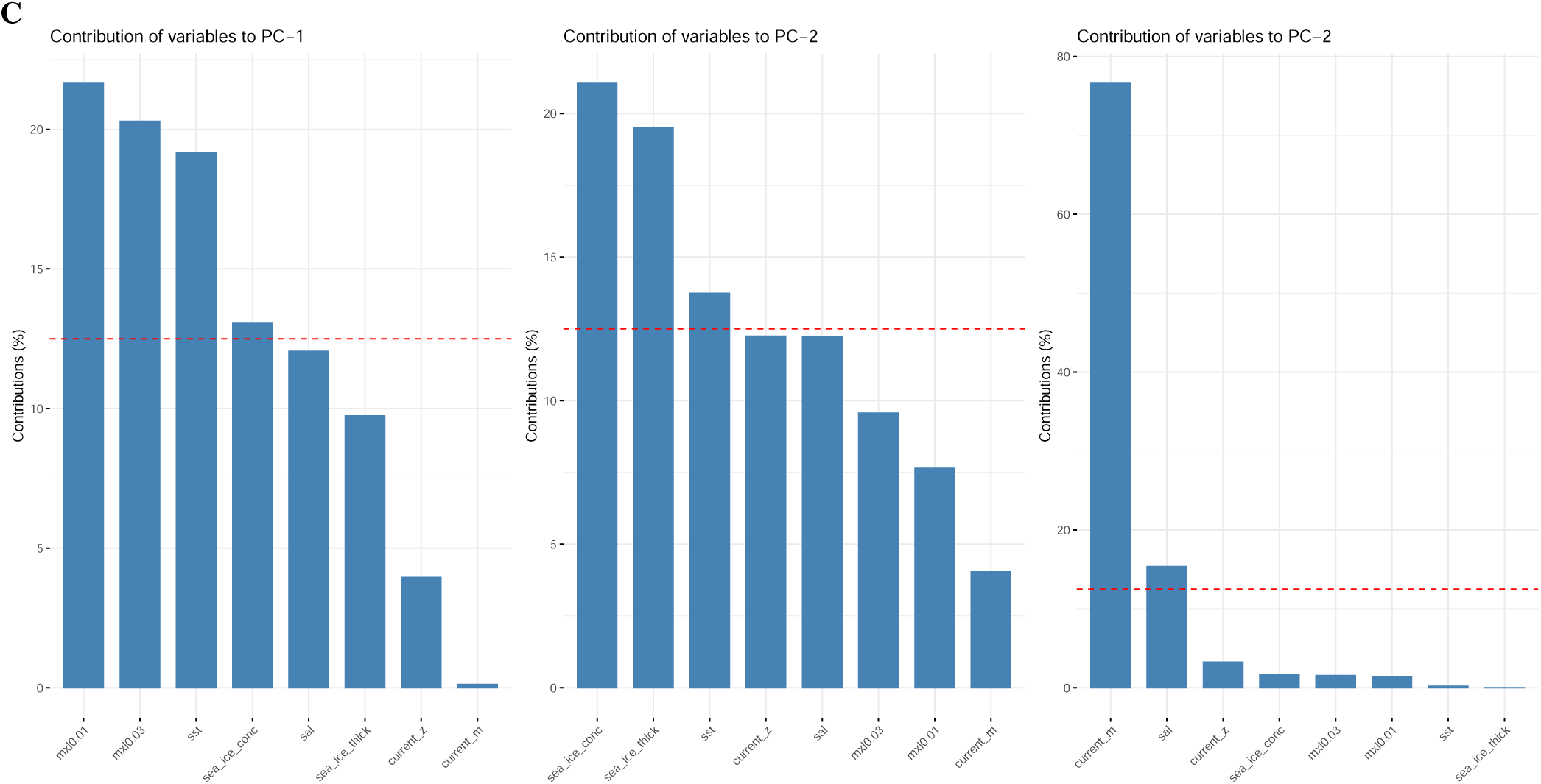
Contributions to the first three principal components from principal component analyses (PCAs) of all eight environmental variables from scenarios (A) Point of Capture–Time of Capture (POC-C), (B) Spawning Ground–Time of Birth (SG-B), and (C) Point of Capture–Time of Birth (POC-B). *sea*_*ice*_*thick*, sea ice thickness; *sea*_*ice*_*conc*, sea ice concentration; *mxl*0.01, mixed-layer depth at 0.01 density; *mxl*0.03, mixed-layer depth at 0.03 density; *sal*, salinity; *sst*, sea surface temperature; *current*_*m*, meridional current velocity; *current*_*z*, zonal current velocity.

**Figure S12.**
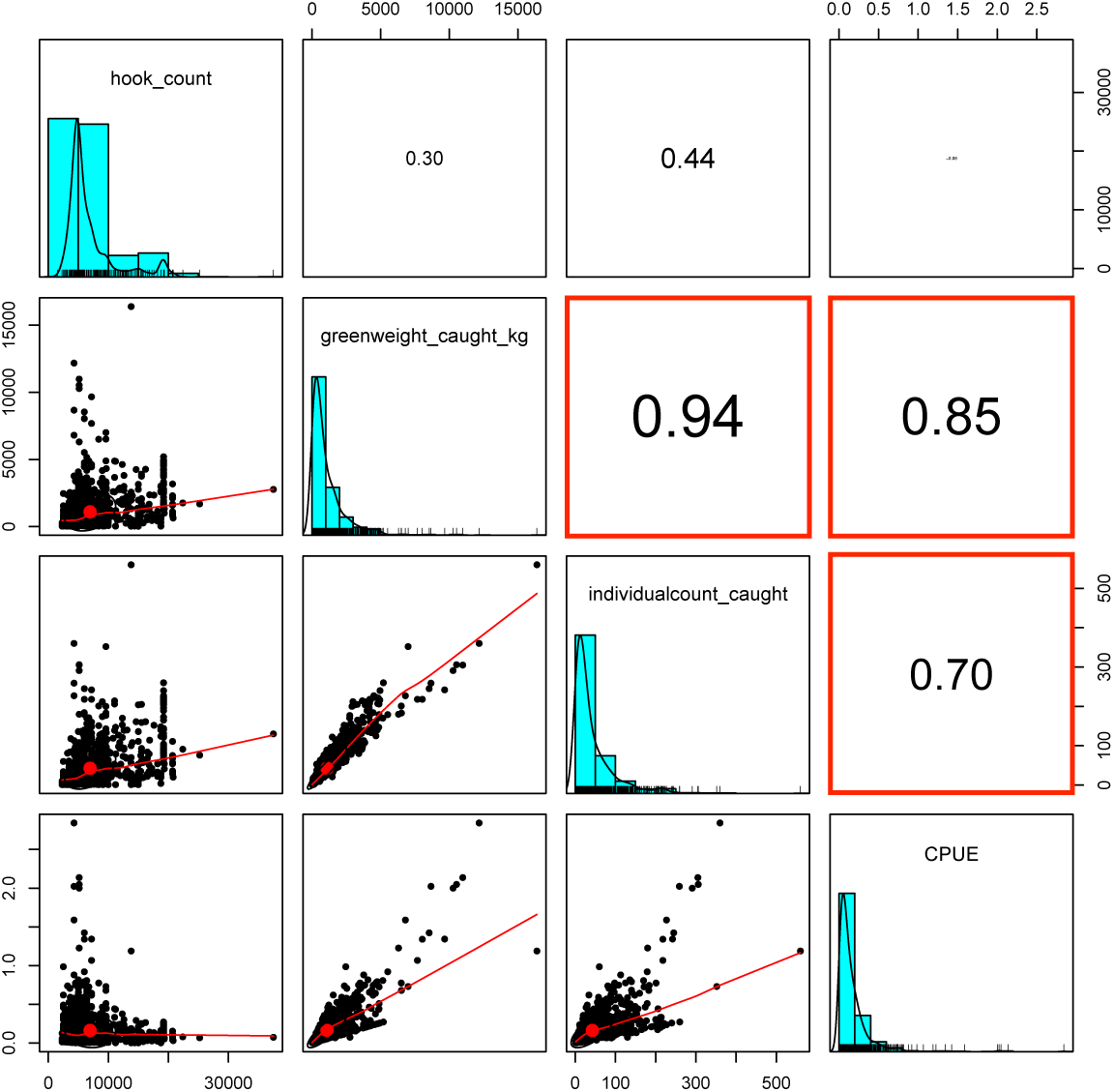
Pairs plots showing collinearity among fishing variable values per haul across all requested years and areas. *hook*_*count*, number of hooks set; *greenweight*_*caught*_*kg*, fish greenweight in kg; *individualcount*_*caught*, number of individuals caught; *CPUE*, catch per unit effort. Boxes with bold red outlines indicate significant collinearity (*r ≥ |*0.7|).

**Figure S13.**
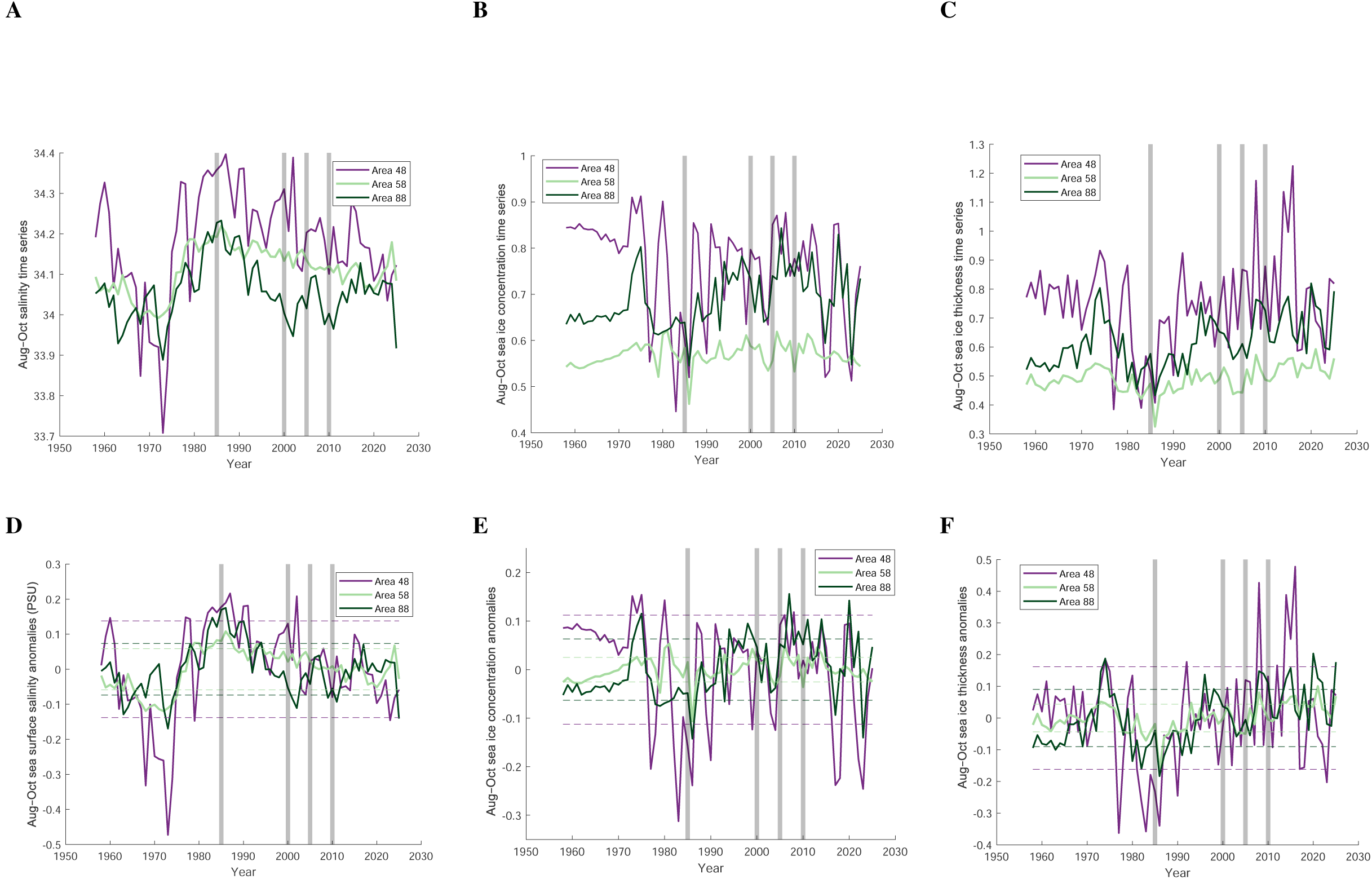
Time series based on raw data (A, B, C) and anomalies (D, E, F) of sea surface salinity (A, D), sea ice concentration (B, E), and sea ice thickness (C, F). Anomalies were derived by subtracting the time series average from each time series point. Raw data from the ORAS5 global ocean reanalysis. Raw data averaged temporally over the estimated *D. mawsoni* egg incubation period (August–October), and spatially over the hypothesized *D. mawsoni* spawning grounds within each CCAMLR area (Figure 1). Dashed lines correspond to one standard deviation from the mean of each of the three time series. The purple line shows data from CCAMLR area 48, the light green line from CCAMLR area 58, and the dark green line from CCAMLR area 88. Shaded gray bars highlight the estimated birth years for *D. mawsoni* used in this study.

